# Signal, Bounds, and Baselines: Principles for Evaluating Virtual Cell Perturbation Models

**DOI:** 10.64898/2026.04.20.719650

**Authors:** Michael Vollenweider, Peter Bühlmann

## Abstract

Foundation models and deep learning systems are increasingly proposed as core components of “virtual cells” capable of forecasting transcriptomic responses to unseen perturbations. Yet rigorous evaluation of perturbation prediction in high-dimensional gene expression space remains challenging, raising concerns of reported performances. Here, we introduce the SBB principles (Signal, Bounds, and Baselines) for evaluating biological perturbation prediction. The Signal pillar introduces diagnostic meta-metrics and promotes differentially expressed gene (DEG)-based weighting or filtering to verify and recover metric sensitivity to biological signal masked by high-dimensional noise. The Bounds pillar provides perturbation-wise metric calibration against empirical reference points, transforming raw metric values into interpretable quantities that clarify the actual scale of model improvements. The Baselines pillar establishes a hierarchy of interpretable linear models that serve as rigorous performance floors. Applying these principles across seven transcriptomic perturbation datasets, including single and double perturbations, we demonstrate that complex deep learning methods, including foundation models, often still fail to meaningfully surpass simple linear baselines, and that substantial room for improvement remains even where they do. These principles provide a critical standard for distinguishing genuine biological signal from statistical artifacts and for guiding more robust model development.

---

Predicting how a cell will respond to an unseen genetic or chemical perturbation is a central goal of computational biology, with direct implications for drug discovery, functional genomics, and cell engineering. High-throughput transcriptomic screens now profile cellular responses to both single (1–4) and combinatorial (5–7) perturbations at single-cell resolution. This data availability, together with the long-term vision of constructing predictive “virtual cells” (8), has driven a large and growing landscape of foundation and domain-specific architectures that aim to predict transcriptomic effects of unseen perturbations (9–21). Whether any of these models genuinely predicts perturbation responses, however, remains an open question.

Despite the rapid surge of new methods, evaluating their predictions within abstract, high-dimensional transcriptomic spaces remains fundamentally challenging (22**?**–27). An ongoing debate has emerged over whether complex deep learning models capture genuine biological insight or merely exploit statistical artifacts that inflate performance under flawed evaluation protocols. Recent systematic benchmarks have revealed what amounts to a widespread “illusion of per-formance”: standard evaluation metrics are undermined by generic systemic biases (28) and by signal dilution in high-dimensional gene expression spaces (29), while state-of-the-art architectures frequently fail to outperform simple predictors such as additive baselines in combinatorial settings and dataset mean baselines for single perturbations (25, 27, 30, 31). These findings call into question not only the effectiveness of current models but also the reliability of the evaluation frameworks used to assess them.

The causes of this evaluation crisis are manifold, and the SBB principles address each directly through three pillars: **Signal, Bounds**, and **Baselines** (Fig. 1). First, commonly used metrics such as MSE and Pearson correlation operate across thousands of genes, the vast majority of which are unaffected by any given perturbation; the resulting signal dilution can render biologically meaningful effects undetectable (29, 32). The **Signal** pillar provides diagnostic meta-metrics to verify that the chosen evaluation metric is sensitive to the biological signal of interest, and promotes signal exposure techniques, such as DEG-based weighting or filtering, to recover signal masked by high-dimensional noise. Second, raw metric values are difficult to interpret in isolation: a small MSE improvement may be negligible or substantial depending on the achievable optimum for a given perturbation. The **Bounds** pillar addresses this by mapping each metric to the interval between empirical upper and lower bounds, transforming them into interpretable quantities that reveal both the magnitude of model improvements and the remaining gap to optimal performance. Third, deep learning models are often compared only against unlearned baselines such as the control mean or additive predictor, overlooking simple learned linear models that may already capture much of the available signal. The **Baselines** pillar establishes a hierarchy of interpretable linear models, from unlearned predictors through learned baselines to analytical oracles, that provide rigorous performance floors and ceilings. Without addressing these interrelated issues, the field risks continued investment in increasingly complex architectures that yield little biological insight. We demonstrate these principles by benchmarking several state-of-the-art methods across seven perturbational transcriptomics datasets (three double perturbation, four single perturbation).

**Fig. 1.**
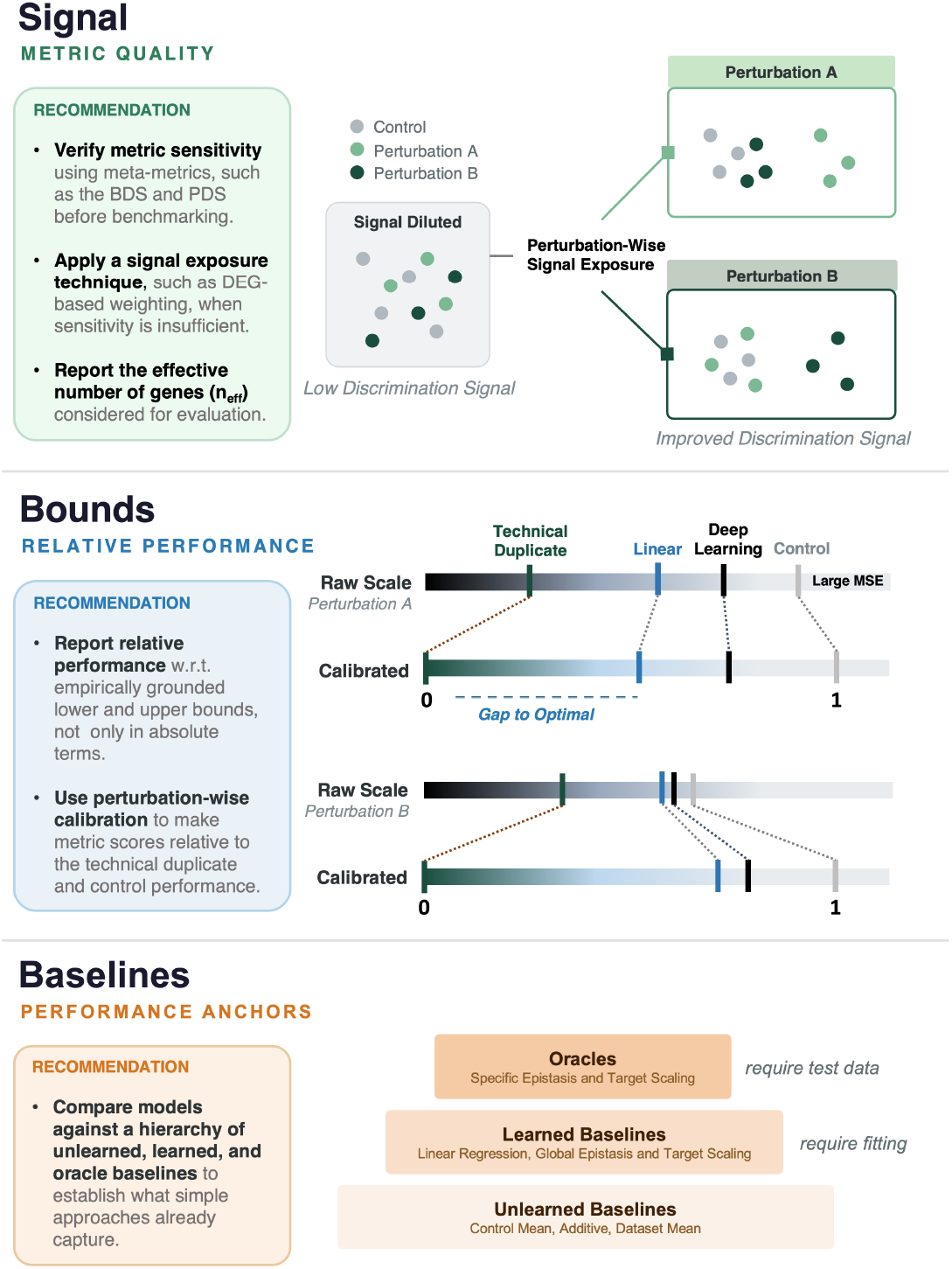
The SBB (Signal, Bounds, and Baselines) principles for perturbation prediction evaluation.

## Results

We evaluate perturbation prediction using three complementary metrics (see Methods): MSE for absolute magnitude, PearsonΔPert for directional alignment of expression changes, and the perturbation discrimination score (PDS) for whether predicted profiles can be correctly matched to their corresponding ground-truth perturbation against all other perturbations. Each metric can be computed in unweighted form across the full gene set or in DEG-weighted form, which concentrates evaluation on genes with substantial perturbation signal. We focus our main results on the advocated weighted variants (wMSE, wPearsonΔPert, wPDS) and additionally report unweighted results for completeness.

To contextualize deep learning performance, we compare against linear baselines of increasing sophistication (see Methods). *Unlearned baselines* require no training and include the control mean, dataset mean, and additive predictor (for combinatorial screens). *Learned baselines* fit simple parametric models on training data, such as a linear regression, a linear epistasis model that captures systematic deviations from additivity through a shared scaling factor and gene-wise offsets, and a target scaling predictor for single perturbation screens, adapted from Wong et al. (33). Finally, *oracles* require access to test data and therefore serve as theoretical upper bounds rather than valid predictors; for example, the specific epistasis oracle estimates combination-specific scaling coefficients from test set perturbation means.

We first compare models against this hierarchy across seven perturbation datasets (Fig. 2), encompassing three combinatorial screens: wessels23 (7), norman19 (5), and replogle20 (6), and four single perturbation screens: frangieh21 (2), adamson16 (1), replogle22k562 (3), and nadig25hpeg2 (4), showing that deep learning models often fail to surpass even simple linear predictors. We then motivate and apply perturbation-wise metric calibration to reveal that even where deep learning models rank favorably, their absolute gains over baselines are small and the gap to oracle performance remains large (Fig. 3). Finally, we analyze the metrics themselves, demonstrating that DEG weighting much better recovers biological signal masked by high-dimensional noise and and we quantify signal dilution through the Boundary Discrimination Score (BDS) and Perturbation Discrimination Score (PDS) meta-metrics (Fig. 4, 5).

**Fig. 2.**
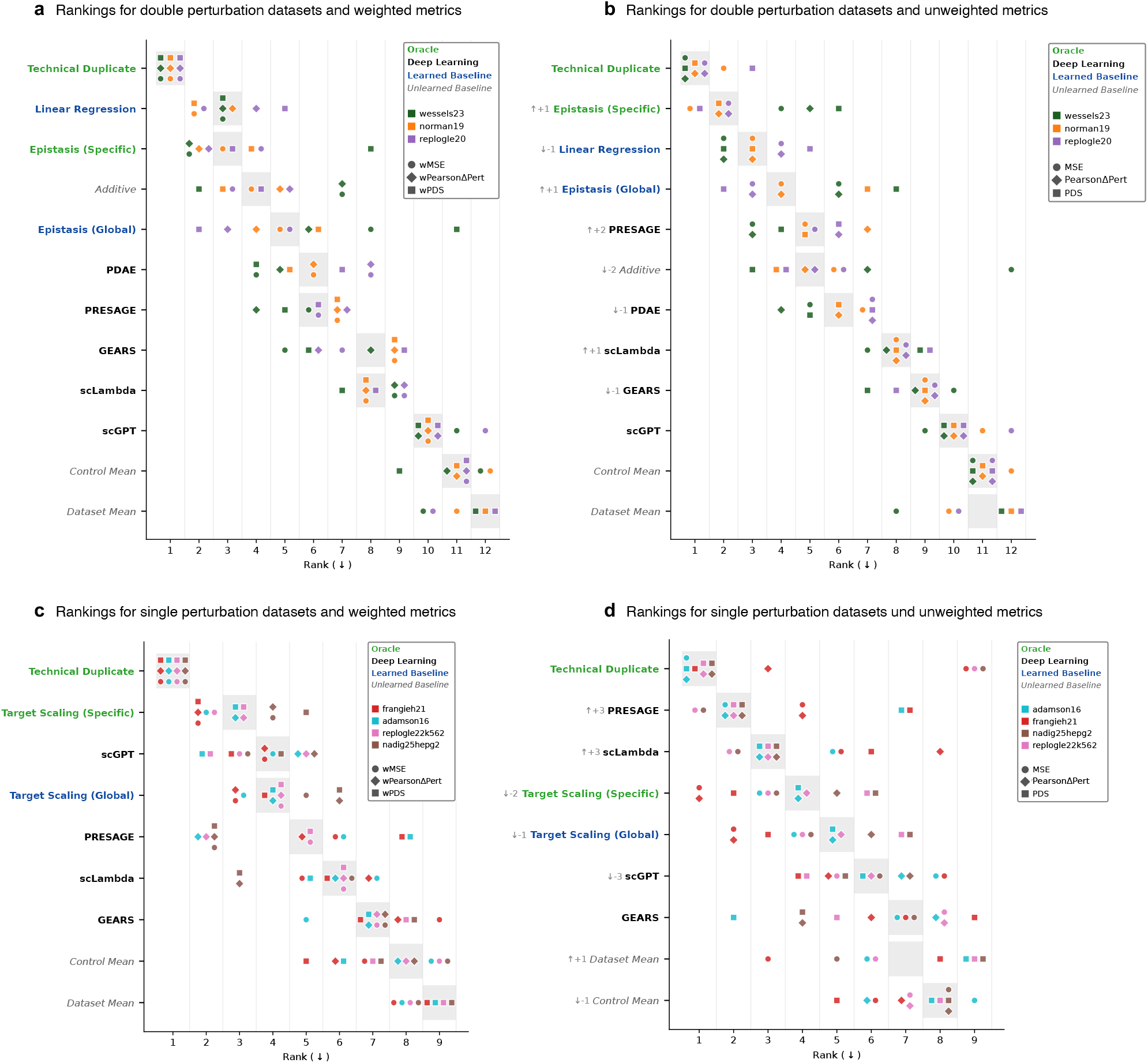
Model rankings across perturbation datasets. **a**, Rankings for double perturbation datasets under weighted metrics (wMSE, wPearsonΔPert, wPDS). **b**, Rankings for double perturbation datasets under unweighted metrics (MSE, PearsonΔPert, PDS). **c**, Rankings for single perturbation datasets under weighted metrics. **d**, Rankings for single perturbation datasets under unweighted metrics. Colors distinguish datasets; shapes distinguish metrics. Models are ordered by median rank across all metric-dataset combinations within each panel. The areas shaded in grey mark the overall median rank across datasets and metrics. Arrows in grey for unweighted metric plots indicate increase or decrease of model rank relative to the ranking for the weighted metrics.

**Fig. 3.**
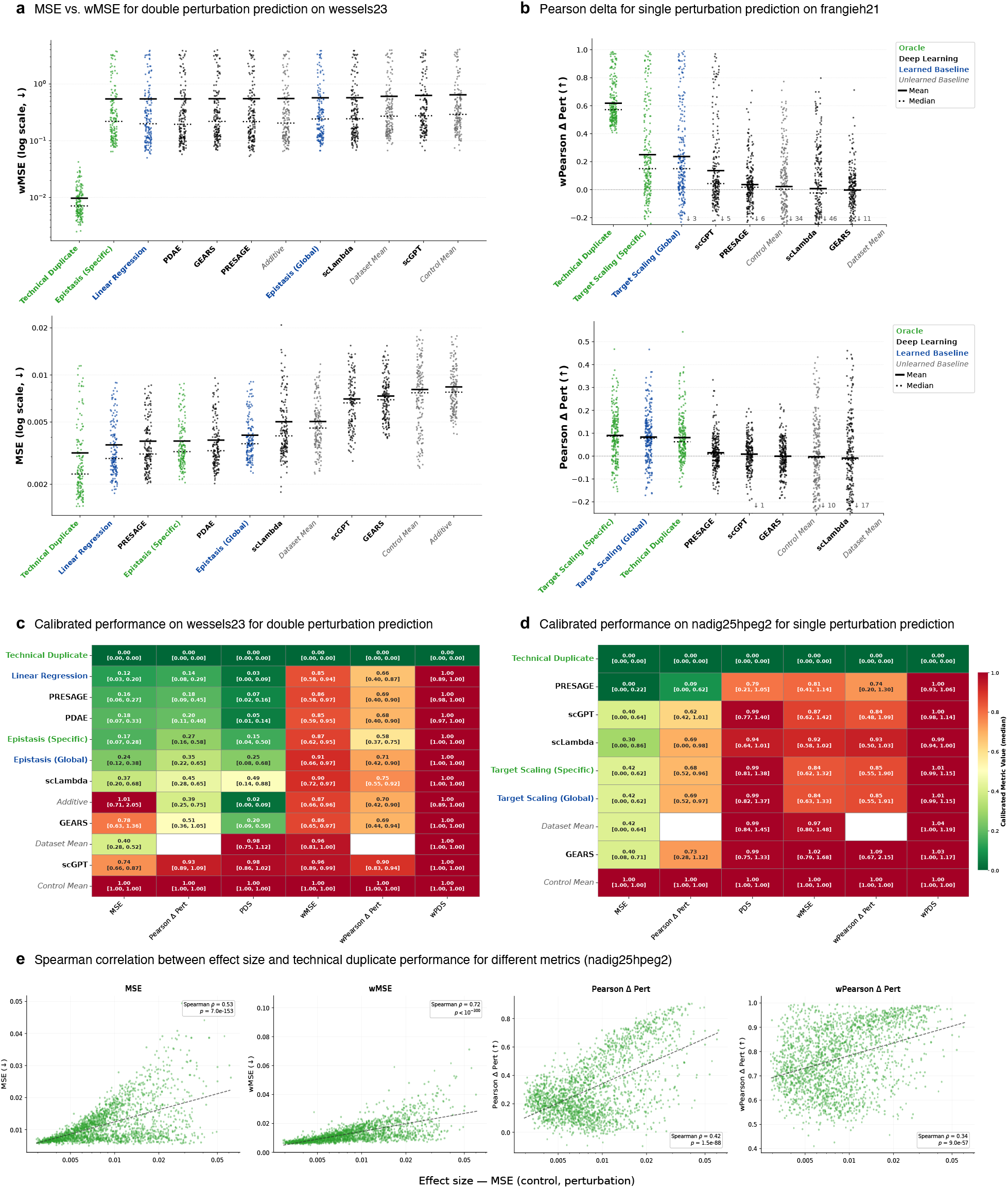
Metric calibration reveals limited model improvement. **a**, MSE versus wMSE for double perturbation prediction on wessels23, showing per-perturbation metric values across models. **b**, PearsonΔPert for single perturbation prediction on frangieh21, comparing weighted and unweighted variants. For both stripplots, models are sorted by mean performance across perturbations **c**, Calibrated performance heatmap on wessels23 for double perturbation prediction; values represent median calibrated scores (0 = technical duplicate, 1 = uninformative control) and brackets indicate the 25th and 75th percentile. **d**, Calibrated performance heatmap on nadig25hpeg2 for single perturbation prediction. For both heatmaps, models and metrics are sorted by mean median performance. **e**, Spearman correlation between perturbation effect size and technical duplicate performance for different metrics on nadig25hpeg2. Calibrated heatmaps for all datasets and further stripplots are provided in Supplementary Note 1& 2.

**Fig. 4.**
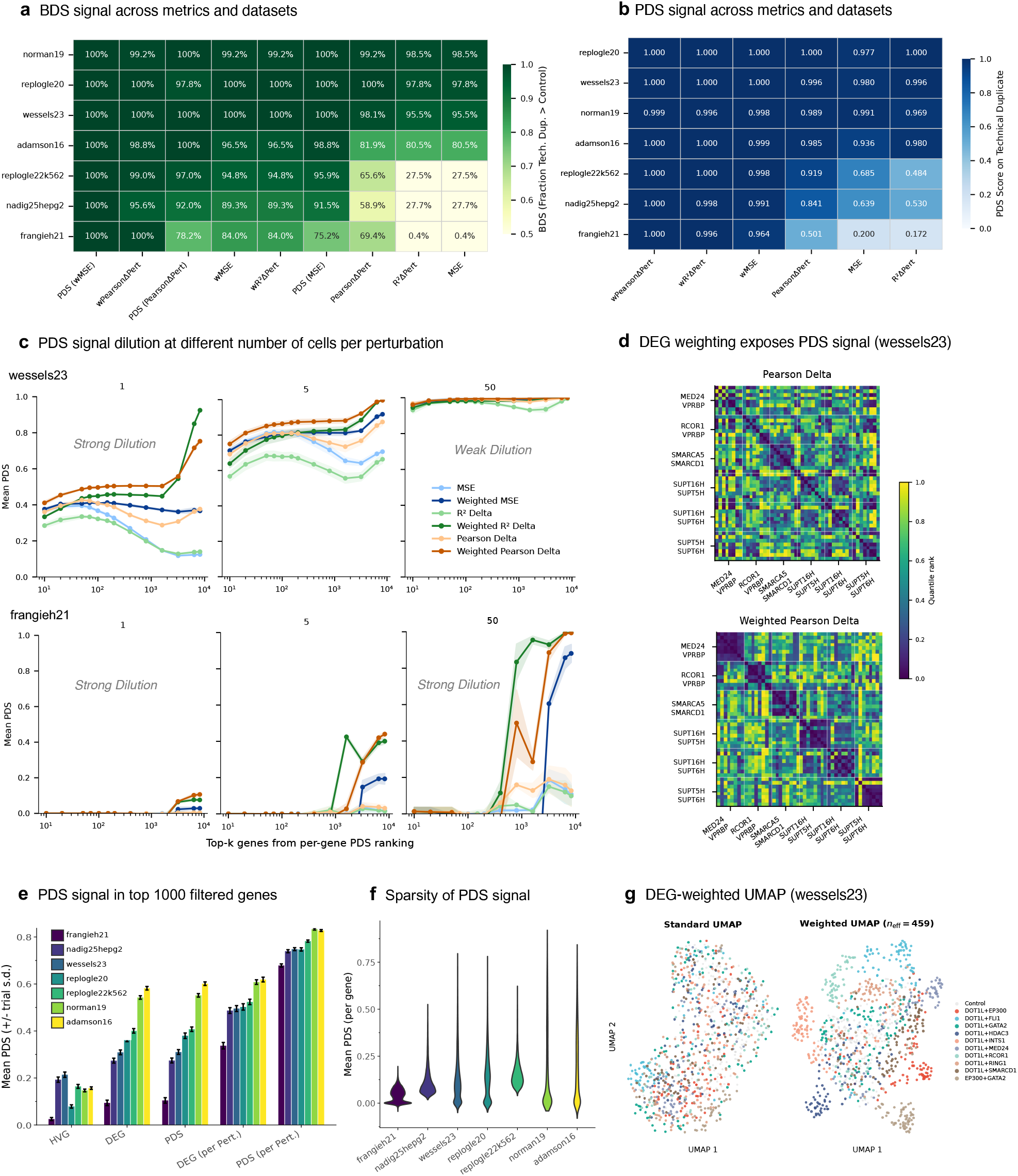
Signal sensitivity of evaluation metrics. **a**, Boundary Discrimination Score (BDS) across metrics and datasets. **b**, Perturbation Discrimination Score (PDS) on technical duplicates across metrics and datasets. It shows the mean over all test folds (2 for double, 5 for single) of the mean PDS across all test perturbations in these folds. **c**, PDS signal dilution at different cell counts per perturbation for wessels23 (top) and frangieh21 (bottom), showing how adding uninformative genes can degrade PDS signal for unweighted metrics. For frangieh21, signal dilution persists even at high cell counts. **d**, Pairwise cell-cell distance matrices for wessels23 under PearsonΔPert versus weighted PearsonΔPert. Color indicates the quantile rank of a cell-cell distance in a given row. DEG weights for the weighted metric are specific to the perturbation of a given row, which makes the weighted distance matrix asymmetric. **e**, PDS signal in top 1000 filtered genes across filtering strategies with ranking according to most Highly Variable Genes (HVG), Differentially Expressed Genes (DEG), and gene-wise PDS signal (PDS), all mean-aggregated across all perturbations, and DEG and gene-wise PDS but perturbation-specific. **f**, Sparsity of gene-wise PDS signal mean-aggregated across perturbations for different datasets. **g**, Standard versus DEG-weighted UMAP of wessels23, illustrating how weighted metrics improve visualization of perturbation heterogeneity. Here, DEG weights are mean-aggregated across perturbations. A more comprehensive comparison of metrics can be found in Supplementary Note 7.

**Fig. 5.**
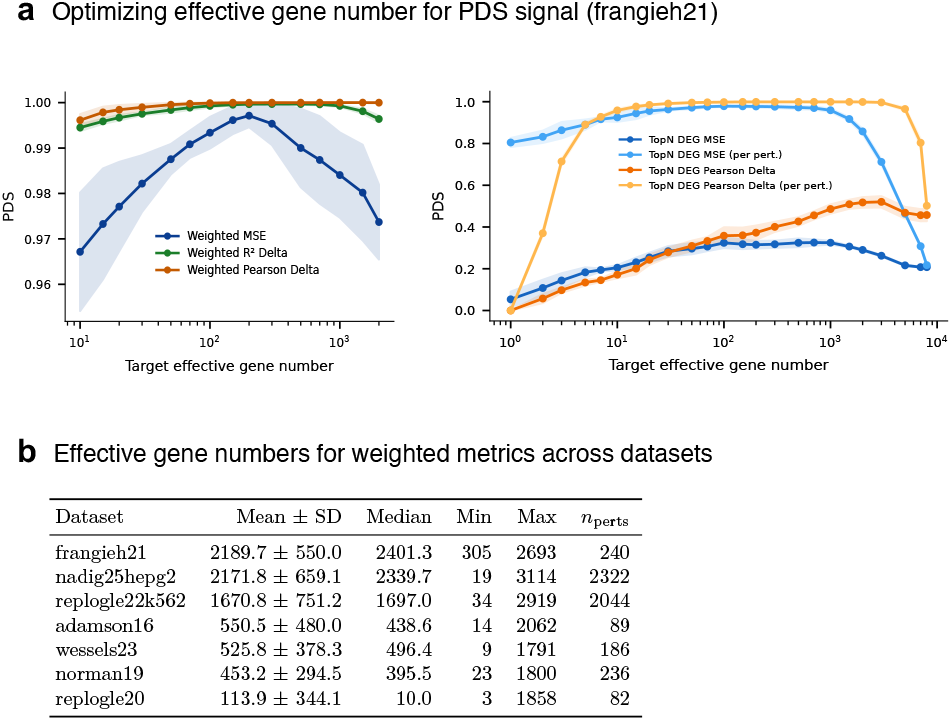
Weighting optimization and effective gene numbers. **a**, Optimizing the effective gene number for PDS signal on frangieh21 for top-*n* metrics and DEG-weighted metrics. For TopN metrics without perturbation-specific gene selection, the weights across all perturbations were mean aggregated for selecting topN genes. **b**, Effective gene number (*n*_eff_) statistics across perturbations for each dataset, reported for the weighting scheme used throughout.

### Simple linear baselines often outperform deep learning models

For double perturbation prediction (Fig. 2a), none of the evaluated deep learning models, including GEARS (11), PRESAGE (21), scLAMBDA (34), PDAE (35), and scGPT (19), outperformed the linear regression baseline across any weighted metric, for all datasets. Further, none of them consistently outperform the unlearned additive and learned epistasis baselines. The result is particularly noteworthy for wessels23, where it has been claimed that models such as PRESAGE consistently surpass the additive baseline (36). While this claim is confirmed under wMSE and wPearsonΔPert, the linear regression baseline clearly outperforms all deep learning models across all three metrics on this dataset. The apparent success of deep learning models thus reflects the weakness of the additive baseline on wessels23 rather than genuine model sophistication. For single perturbation prediction (Fig. 2c), the target scaling baseline frequently matches or exceeds deep learning model performance, particularly on the smaller datasets frangieh21 and adamson16. Under unweighted metrics (Fig. 2b,d), deep learning models gain relative standing in both settings, a pattern we attribute to signal dilution masking their failures on biologically relevant genes. These findings echo mounting evidence from recent systematic benchmarking efforts (25, 27, 30, 31) that complex architectures routinely fail to surpass simple predictors. We note, however, that even where deep learning models outperform simple baselines in rankings, as on nadig25hpeg2, the calibrated analysis below will reveal that models remain far from the achievable optimum.

### Learned baselines and oracles illuminate performance gaps

For wessels23, the global epistasis baseline substantially outperforms the additive predictor (Fig. 3a) on unweighted MSE, indicating that this dataset exhibits pronounced non-additive structure, potentially attributable to systemic buffering, that a purely additive model cannot capture. Notably, this model captures only genome-shared epistasis: a single scaling factor applied to the sum of single-perturbation effects, plus an offset, applied uniformly across all genes, without any pathway- or gene-specific interaction terms. Yet it reaches performance competitive with state-of-the-art deep learning models. An analogous pattern emerges for single perturbation prediction, where the target scaling baseline, which simply sets the target gene’s expression to a learned penetrance fraction while leaving all other genes at the dataset mean, frequently matches or exceeds deep learning models on frangieh21 and adamson16 (Fig. 2c, 3b). That such a minimal, single-parameter intervention on a single gene can rival complex architectures underscores how little signal current models extract beyond what is trivially available from the perturbation’s identity alone. This carries an important practical implication: one should consider not only unlearned baselines but also simple learned baselines, as the gap between unlearned and learned linear models can be substantial and reveals how much signal is accessible to models that lack biological priors entirely.

### Metric calibration reveals substantial room for model improvement

While ranking-based comparisons establish relative model orderings, they obscure the absolute magnitude of performance gains. To quantify the practical significance of observed improvements, we introduce perturbation-wise metric calibration. Throughout, we refer to two empirical reference points: an upper bound (or positive baseline) derived from technical duplicate agreement and a lower bound (or negative baseline) derived from uninformative con-trol samples. Calibration maps each model’s raw metric value to the interval defined by these bounds: a score of 0 corresponds to technical duplicate performance (the empirical optimum), while a score of 1 corresponds to the uninformative baseline; values exceeding 1 indicate performance worse than the negative control (see Methods). An additional motivation for perturbation-wise calibration is that the relationship between perturbation effect size and raw metric performance is metric-dependent: larger effect sizes correlate with worse MSE but better PearsonΔPert (Fig. 3e), so that different perturbations dominate the aggregate score depending on the metric. Calibration normalizes these scale differences, enabling fair comparison across perturbations, datasets, and metrics.

Applying this calibration across datasets reveals a consistent pattern: even where deep learning models surpass simple baselines in rankings, the calibrated improvement is modest relative to the achievable optimum. On wessels23 (Fig. 3c), models such as PDAE, GEARS, and PRESAGE rank above the additive baseline for weighted metrics, yet their calibrated scores remain far from the technical duplicate bound, indicating that most of the achievable performance gap remains unclosed. On nadig25hpeg2 (Fig. 3d), a similar picture emerges: models occupy only a narrow band between the uninformative baseline and the technical duplicate under calibrated evaluation. The calibrated view also exposes metric-dependent blind spots. PRESAGE on nadig25hpeg2 achieves strong calibrated scores under unweighted metrics (median calibrated MSE: 0.00; PearsonΔPert: 0.09) yet performs poorly under weighted variants (wMSE: 0.81; wPearsonΔPert: 0.76), indicating that its predictions are accurate on genes unaffected by the perturbation but fail to capture the biologically relevant expression changes. Calibrated heatmaps for all datasets are provided in Supplementary Note 1 and metric-wise stripplots for wessels23 and nadig25hpeg2 in Supplementary Note 2.

Notably, our calibrated metrics trivially saturate the Dynamic Range Fraction (DRF) introduced by Miller et al. (36): because the calibrated score is by construction a linear transformation relative to the bounds, the DRF evaluates to exactly 1 for every valid perturbation, questioning the suitability of the DRF as a standalone meta-metric.

### DEG-based weighting recovers sensitivity of metrics to biological signal

Before benchmarking predictive models, it is essential to verify that the chosen evaluation metric possesses sufficient sensitivity to detect genuine biological signal within the ground-truth data. We assess this using two diagnostic meta-metrics computed exclusively from experimental observations (see Methods). The Bound Discrimination Score (BDS) measures the fraction of perturbations for which a metric correctly ranks a technical replicate closer than an uninformative control, assessing metric sensitivity to perturbation effects against background noise. The Perturbation Discrimination Score (PDS), repurposed here as a meta-metric by computing it exclusively on technical duplicate samples, measures whether the chosen dissimilarity can resolve one perturbation from another. Together, these diagnostics provide a straightforward check of metric adequacy prior to any model benchmarking. The BDS analysis (Fig. 4a) reveals that standard unweighted MSE fails to reliably discriminate technical replicates from controls across several datasets, with BDS values as low as 27.5% for re-plogle22k562 and nadig25hpeg2, and 0.4% for frangieh21. By contrast, DEG-weighted MSE (wMSE) recovers a moderate or good BDS across all datasets (84 – 100%). Unweighted PearsonΔPert already demonstrates greater resilience than MSE, consistent with its invariance to global scaling, though weighting further improves its fidelity. The rank-based PDS with weighted MSE as the underlying dissimilarity achieves perfect BDS across all datasets, confirming the advantage of rank-based evaluation (30, 32). Notably, *R*^2^ΔPert and MSE yield identical BDS values, as their calibrated forms are algebraically equivalent (Supplementary Note 4). The BDS diagnostic also serves as a prerequisite for meaningful calibration: when it confirms that the upper bound reliably separates from the lower bound, the calibration framework produces interpretable performance scales; when it does not, calibrated scores should not be relied upon.

The PDS analysis (Fig. 4b) empirically substantiates concerns raised by Liu et al. (32) regarding the limited sensitivity of MSE for perturbation discrimination in high-dimensional noisy data. For the large-scale single-perturbation datasets replogle22k562 and nadig25hpeg2, unweighted MSE yields low PDS values (the PDS is scaled to the range [−1, 1], where 0 indicates random ordering; see Methods), demonstrating a fundamental inability to fully resolve individual perturbation signatures. The smaller frangieh21 dataset is similarly afflicted. Correlation-based metrics such as PearsonΔPert perform substantially better (PDS scores of 0.919 and 0.841 for replogle22k562 and nadig25hpeg2), though frangieh21 remains challenging even under PearsonΔPert (0.501). Crucially, DEG-based weighting restores strong perturbation discrimination across all metrics and datasets. We do not recommend abandoning unweighted metrics entirely: for datasets where both weighted and unweighted correlation-based metrics achieve good discrimination (e.g., wessels23, norman19, replogle20, adamson16), unweighted metrics evaluate the complementary aspect of accuracy across the full transcriptome. However, for datasets such as frangieh21, replogle22k562, and nadig25hpeg2, unweighted MSE-based evaluation is fundamentally flawed. Even when unweighted sensitivity is confirmed, weighted metrics remain essential for assessing performance specifically on the genes driving the perturbation phenotype. These meta-metrics provide a rapid diagnostic prior to any model benchmarking. To formalize this assessment, we additionally construct distance-based permutation tests for both the BDS and PDS (Supplementary Note 5) to assess the robustness of these findings.

### High-dimensional noise dilutes biological signals in standard metrics

To elucidate why DEG-based weighting is so effective, we investigate signal dilution in high-dimensional transcriptomic spaces. Supplementary Note 3 formalizes how distance concentration in high dimensions causes sparse perturbation signatures to become indistinguishable from noise under standard metrics, and shows that two complementary strategies, pseudobulking (reducing the noise floor) and gene weighting (increasing the effective signal fraction), systematically restore metric sensitivity.

Perturbation-specific signal is empirically sparse: the genewise PDS analysis (Fig. 4e,f) demonstrates that only a small fraction of genes carry distinguishing information for any given perturbation when considered independently, and that the informative gene sets differ substantially across perturbations. This heterogeneity is evident in Fig. 4e: globally selected top genes yield lower signal than genes selected individually per perturbation. Consequently, evaluation across the full transcriptome without weighting allows uninformative genes to dominate the metric.

The signal dilution effect is further characterized by studying how PDS on technical duplicates varies with gene count and cell count (Fig. 4c). For wessels23, unweighted MSE exhibits severe signal dilution at low cell counts, mitigated by increasing pseudobulking. Frangieh21 presents a more severe case: signal dilution persists even at high cell counts, confirming that pseudobulking alone cannot rescue metric sensitivity for all datasets. Weighted metrics maintain high PDS across conditions for both datasets. The practical consequence is directly visualized in the pairwise distance matrices of wessels23 (Fig. 4d) and UMAP embeddings (Fig. 4g): unweighted PearsonΔPert produces more homogeneous distance structure and diffuse embeddings, while weighted PearsonΔPert reveals clear block-diagonal structure and perturbation groupings.

Because signal exposure techniques alter the effective evaluation dimensionality, we consistently report the effective gene number *n*_eff_ (see Methods; Fig. 5b) alongside all weighted evaluations. The choice of weighting scheme and the resulting *n*_eff_ can itself be optimized; for example, for top-*n* DEG filtering (Fig. 5a), enabling a data-driven choice rather than the arbitrary selection of 20 or 100 genes common in current practice. A comprehensive comparison of DEG-weighted, DEG-filtered, and distributional metrics is provided in Supplementary Note 7.

## Discussion

The SBB principles address a critical gap in the perturbation prediction field: the lack of a unified evaluation framework that diagnoses signal dilution, provides interpretable performance scales, and anchors comparisons to rigorous baselines. Applying these principles across seven transcriptomic datasets, we find that current evaluation practices systematically overestimate model performance and that complex deep learning architectures frequently fail to surpass simple linear baselines when assessed on informative metrics. We organize our discussion around the three pillars of the SBB principles. The resulting recommendations can be summarized as follows:

**Summary of Recommendations**

**Signal:** Verify metric sensitivity using meta-metrics, such as the BDS and PDS, before benchmarking. When sensitivity is insufficient, apply DEG-based weighting, top-n DEG filtering, or another signal exposure technique and report the effective gene number.

**Bounds:** Report performance relative to empirically grounded reference points, not only in absolute terms. For instance, use perturbation-wise calibrated scores relative to the technical duplicate (upper) and uninformative control (lower) bounds.

**Baselines:** Compare models against a hierarchy of unlearned, learned, and oracle baselines to establish what simple approaches already capture. If you include prior biological information in your model, consider using simple learned baselines that also incorporate it.

The **Signal** pillar requires verification that the chosen evaluation metric possesses adequate sensitivity to the biological signal of interest prior to model benchmarking. The BDS and PDS meta-metrics provide simple diagnostics for this purpose and can be extended to permutation tests for formal significance assessment (Supplementary Note 5). When they reveal insufficient discrimination, the metric should not be trusted for model comparison. DEG-based weighting or filtering, or pathway-focused evaluation (37, 38) should be applied instead, with the effective gene number *n*_eff_ consistently reported to ensure that evaluation does not collapse onto a trivially small gene subset. The weighting scheme and *n*_eff_ can themselves be optimized (Fig. 5a), and we encourage future work to investigate what constitutes an optimal weighting scheme (Supplementary Note 7). While the precise technique appears to matter less for recovering the signal, it determines what aspect of the perturbation response is evaluated, so it should be chosen with a clear sense of what is being measured. Beyond evaluation, the signal characterization provided by the BDS and PDS is directly relevant to training objectives: a model trained with an unweighted loss across the full transcriptome may expend capacity predicting noise genes rather than learning perturbation-specific effects. Weighted training objectives (29) and representation learning in signal-enriched subspaces represent promising directions for aligning training with evaluation. In principle, the BDS and PDS also enable direct, model-agnostic comparison of preprocessing pipelines by quantifying how well normalization, transformation, and gene selection choices preserve perturbation signal and discriminability.

The **Bounds** pillar requires that performance be reported not only in absolute terms but also relative to empirically grounded reference points. Rankings alone obscure whether a top-performing model has closed most of the gap to optimal performance or has barely moved beyond the uninformative baseline. Perturbation-wise metric calibration resolves this ambiguity by mapping raw values to the interval between an empirical upper bound (technical duplicate) and a lower bound (uninformative control), exposing both the magnitude of improvements and the remaining gap. Calibration also treats all perturbations on equal footing regardless of effect size, preventing evaluation from being dominated by a subset of perturbations. By construction, calibrated scores also maximize the Dynamic Range Fraction (DRF) (36), yielding a DRF of exactly 1 for every valid perturbation.

The **Baselines** pillar requires that model comparisons be anchored against a hierarchy of baselines of increasing sophistication, not only unlearned predictors such as the control mean or additive baseline. The standard practice of comparing deep learning models only against such uninformative references sets the bar too low, since it overlooks simple learned models that may already capture much of the available signal. We propose a hierarchy spanning unlearned baselines, learned linear models, and oracle variants that provide theoretical upper bounds. This hierarchy serves a dual purpose beyond providing performance floors. Studying its structure yields direct biological insight: the superiority of the global epistasis model over the additive baseline on wessels23, for instance, reveals non-additive structure consistent with systemic buffering (5, 39, 40). Such insights can inform model design, as exemplified by architectures that explicitly build upon additive baselines (41). We recommend that practitioners systematically evaluate this baseline hierarchy, which provides a clear picture of what simple approaches already capture and where genuine room for improvement exists. The oracle hierarchy also enables finer error decomposition: subtracting the specific epistasis oracle from the ground truth isolates gene-specific epistasis as an evaluation target, complementing the additive-residual approach of Ahlmann-Eltze et al. (31).

Beyond improving how we evaluate perturbation prediction, our analyses raise a more fundamental question: whether the task as currently framed is achievable from transcriptomic data alone. The sparsity and heterogeneity of perturbation-specific gene sets (Fig. 4d,e) imply that the effective training sample size for any given test perturbation can be dramatically smaller than the nominal number of training perturbations, since only perturbations with overlapping active gene sets provide relevant learning signal. Moreover, even when a model does appear to predict an unseen perturbation well, it may simply be interpolating within a cluster of functionally similar perturbations rather than generalizing across mechanistically distinct ones. The mathematical conditions under which such out-of-distribution extrapolation is possible remain poorly understood, and current methods rarely make their structural assumptions explicit. The challenge may also run deeper than data scarcity. Perturbation effects propagate through chromatin remodeling, post-transcriptional regulation, and protein-protein interactions not directly observed in steady-state mRNA measurements, and mRNA levels can explain only roughly 10% of protein abundance variance (42). A perturbation whose mechanism is not reflected in the training transcriptomes cannot be predicted from transcriptomic patterns alone (43). Whether unimodal transcriptomics contains sufficient information for reliable extrapolation across mechanistically distinct perturbations remains unclear. Multimodal measurements integrating proteomics (44), chromatin accessibility (45), or spatial readouts (46) may be necessary for qualitatively different information about the regulatory layers that mediate perturbation responses.

Several limitations of the present work point to important directions for future research. Our implementation of each SBB pillar represents an initial instantiation designed to establish the conceptual framework; each can be substantially extended. In particular, the DEG-based weighting scheme we employ is one of several plausible choices, and the resulting weight distributions vary considerably across datasets (as reflected in the range of effective gene numbers *n*_eff_ reported in Fig. 5b). Once sufficient signal is guaranteed, the choice of weighting scheme becomes less a statistical question than a scientific one about what aspect of the perturbation response one wants to predict well, and a systematic comparison of alternatives should therefore involve domain experts who can assess whether the prioritized genes align with the biology of interest. Our benchmarking spans seven datasets and a representative subset of methods; broader evaluation across additional cell types, perturbation technologies, and emerging modalities will be necessary to establish full generality. Furthermore, our analysis focuses on mean expression predictions; distributional methods and baselines warrant dedicated evaluation under the SBB principles. Finally, the learned baselines themselves are intentionally simple and could be enriched while remaining interpretable, for instance by incorporating pathway annotations or regulatory priors.

## Methods

### A. Notation and setup

We consider perturbation prediction in the setting of single-cell transcriptomics. Let *n*_genes_ denote the number of measured genes. For a perturbation condition *p*, we observe a set of single cells 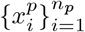, where each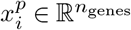 represents a preprocessed and log-transformed gene expression vector. We denote the control (unperturbed) mean expression by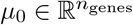, the perturbation mean by 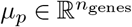, and the mean perturbation effect (log-fold change) by Δ_*p*_ = *µ*_*p* −_ *µ*_0_. For combinatorial perturbation conditions *p*+*q*, we write *µ*_*pq*_ for the observed combination mean. Throughout, 𝒫 denotes the set of all perturbation conditions excluding the control, with | 𝒫 | = *n*_perts_.

### B. Baselines

We consider a suite of linear models to establish rigorous performance floors and ceilings. These are organized into three categories: standard unlearned baselines that require no training, learned baselines that are fitted on training data, and oracle models that additionally require access to ground-truth test data.

#### Unlearned baselines

The *control mean baseline* predicts 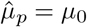for all conditions. The *dataset mean baseline* predicts 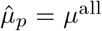, where *µ*^all^ is the global mean expression across all training cells, providing a negative baseline that avoids control bias (28). The *additive baseline* predicts combinatorial effects as the sum of individual perturbation effects: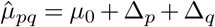.

#### Global epistasis model (learned baseline)

This model assumes that double perturbation effects are scaled, additive versions of the single perturbation profiles, with a shared systemic offset. We model the combination mean as

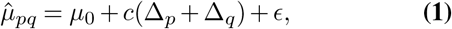

where *c* ∈ ℝ is a scaling factor and 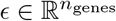 is a gene-wise offset, both shared across all perturbation conditions. Then, *c* and *ϵ* are estimated jointly across all training and validation combinations via ordinary least squares (OLS). We term this the “global” epistasis model because its parameters are shared across all perturbation combinations, allowing it to be fit on training data alone. This model captures uniform epistatic effects such as synergy and buffering, that is, deviations from strict additivity that are consistent in magnitude across all genes, phenomena known in genetic interaction studies (40).

#### Specific epistasis model (oracle)

This variant extends the global epistasis model by fitting combination-specific scaling coefficients:

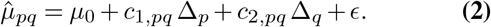

The pair-specific scalars (*c*_1,*pq*_, *c*_2,*pq*_) for all combinations and the shared offset *ϵ* are estimated simultaneously via a single joint OLS fit. This formulation is conceptually related to the combination-specific epistasis analysis of Replogle et al. (6), though our model differs in an important respect: we include a shared gene-wise offset *ϵ* to capture systemic effects common to all perturbation combinations, as identified by (28). The pair-specific scalars explicitly quantify the variance explained by combination-specific interactions. Because computing the pair-specific coefficients requires access to the ground-truth combination mean *µ*_*pq*_ at test time, this model functions as a theoretical upper bound for linear additive models rather than a valid predictive baseline.

#### Linear regression (learned baseline)

We construct a perturbation vocabulary 𝒫 and encode each condition with a binary indicator vector *z* ∈ {0, 1}^|𝒫|^: the control maps to the zero vector, a single perturbation *p* has a one at index *p*, and a combination *p*+*q* has ones at both indices *p* and *q*. We fit a multi-output linear model on perturbation means:

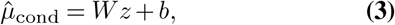

where 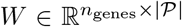 and 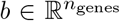. The model is trained by minimizing the *L*_2_ (MSE) loss. This baseline was introduced in (35) and used to evaluate on simulated data.

#### Target scaling baseline (learned baseline & oracle)

Designed for single perturbations, this baseline adapts the methodology of Wong et al. (33), who showed that simply setting the target gene expression to zero for gene knockouts or doubling for gene activations (as in norman19) yields predictions that often exceed deep learning models. Rather than assuming complete knockout or activation, our adaptation learns a scaling factor that captures the dataset’s average penetrance.

Let 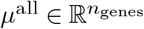denote the global mean expression across all training and validation cells. Each perturbation condition *p* ∈ 𝒫_train_ targets a known subset of genes *T*_*p*_ ⊆ {1,…, *n*_genes_}. We define a masked reference vector 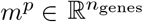that isolates the target genes:

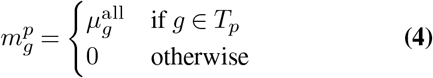

The baseline models the mean perturbed profile as a single-parameter shift applied to the target genes:

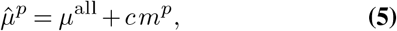

where *c* ∈ ℝ is a scalar penetrance coefficient shared across all perturbations. The coefficient *c* is estimated by ordinary least squares over the training perturbations:

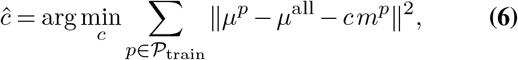

where *µ*^*p*^ is the observed mean expression for perturbation *p*. Taking the derivative and setting it to zero gives the closed-form solution:

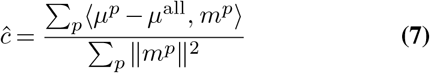

For a test perturbation *q* with target genes *T*_*q*_, the prediction is 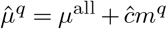, which leaves all non-target genes at *µ*^all^ and scales each target gene by (1 + *ĉ*). The *global* variant estimates *c* from the training set only, while the *specific* (oracle) variant directly sets the target gene to its ground-truth expression value, with all other genes remaining at *µ*^all^.

### C. Evaluation metrics

We organize evaluation metrics into three categories, each capturing a complementary aspect of prediction quality: absolute error (MSE), directional alignment (PearsonΔPert), and distinctness (PDS). All three operate on point estimates of the perturbation mean. Each metric can be computed in unweighted form across the full gene set or in weighted form using DEG-based gene weights (see Signal exposure techniques below). Distributional metrics for comparing single-cell populations are discussed in Supplementary Note 7.

#### Mean squared error (MSE)

Given a predicted perturbation mean 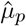 and observed mean *µ*_*p*_, the MSE quantifies absolute prediction error:

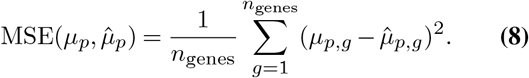

#### PearsonΔPert

Introduced in (11), this metric captures directional alignment by computing the Pearson correlation between the predicted and observed expression changes relative to a negative baseline *µ*^−^:

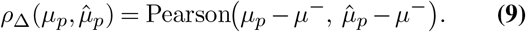

While the control mean *µ*_0_ was originally proposed as the negative baseline (11, 47), Viñas-Torné et al. (28) showed that this choice is susceptible to control bias. Following Miller et al. (36), we use the mean over all training perturbations. We compare different negative baselines as references for the Pearson correlation in Supplementary Note 7. The *R*^2^ Delta Perturbation metric 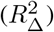 is defined analogously using *R*^2^ instead of Pearson correlation.

#### Weighted point-estimate metrics

To focus evaluation on genes exhibiting significant expression changes, we extend each metric with perturbation-specific DEG-based weights 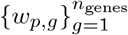 (see Signal exposure techniques). The *weighted MSE* (wMSE) is defined as

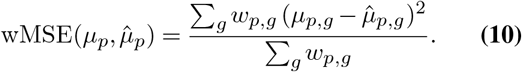

The *weighted Pearson Delta* (wPearsonΔPert) computes a weighted Pearson correlation on the expression changes 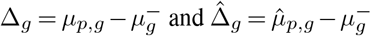. Defining the weighted covariance

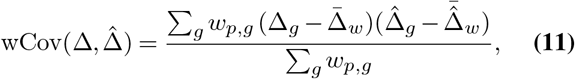

Where 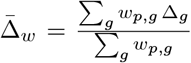 and 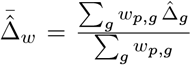 are the weighted means, the metric is

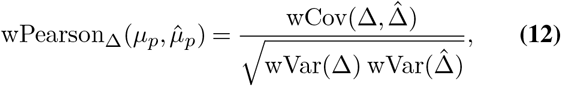

with wVar(Δ) = wCov(Δ, Δ) and analogously for 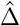. To our knowledge, we are the first to apply continuous gene weighting to the Delta Pearson metric.

#### Perturbation Discrimination Score (PDS)

The PDS quantifies how distinguishable predicted perturbation effects are from one another, and plays multiple roles in our framework. We use it in three distinct modes: (i) as a *meta-metric* for evaluating ground-truth signal strength, computed on technical duplicates (see Evaluating metric signal below); (ii) in a *gene-wise* form, where a separate PDS is computed for each gene individually (using a one-dimensional dissimilarity) to assess the sparsity of perturbation-specific signal; and (iii) as an *evaluation metric* for model predictions, assessing whether predicted perturbation profiles are correctly discriminated from other conditions. In all cases, the underlying computation is identical and parametrized by the choice of dissimilarity measure *d*(·, ·).

Each perturbation *p* is represented by two disjoint sets of cells: a *query* set *Q*_*p*_ and a *reference* set *R*_*p*_. The score measures whether *Q*_*p*_ is closer to its own reference *R*_*p*_ than to the references *R*_*q*_ of other perturbations. Specifically, we compute a within-perturbation set dissimilarity *d*(*Q*_*p*_, *R*_*p*_) and cross-perturbation set dissimilarities *d*(*Q*_*p*_, *R*_*q*_) for all *q* ≠*p*, and evaluate the fraction of cross-perturbation distances that exceed the within-perturbation distance:

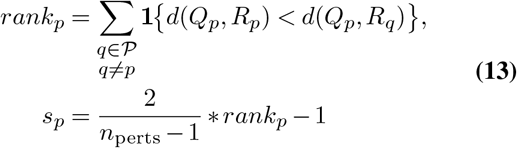

with ties broken uniformly at random. When the dissimilarity *d* is a weighted metric (e.g., wMSE or wPearsonΔPert), the weights used are always those of the query perturbation *p*, ensuring that each perturbation is evaluated on the genes where it induces the strongest signal. The overall PDS is the average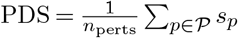. A PDS of 1.0 indicates perfect discrimination, 0 indicates random ordering, and −1.0 indicates systematically incorrect ordering. We note that we report the PDS on the rescaled [−1, 1] interval, rather than as a normalized rank in the range [0, 1], to emphasize that the lowest achievable score indicates a systematic reversal of ordering.

When used as an evaluation metric, *Q*_*p*_ is the model’s predicted profile for perturbation *p* and *R*_*p*_ is the ground-truth profile, so the score measures whether predictions land closer to their corresponding ground truth than to the ground truth of other perturbations. When used as a meta-metric for ground-truth signal quality, the ground-truth samples for each perturbation are partitioned into two equally sized bags, which serve as *Q*_*p*_ and *R*_*p*_ respectively; the resulting score quantifies whether a perturbation’s technical replicates are more similar to each other than to replicates of other perturbations.

The aggregated PDS can be formally interpreted as a rescaled area under the ROC curve (AUROC) for a distance-based binary classification task (Supplementary Note 6).

The PDS has been widely adopted in single-cell perturbation analysis under various names, including retrieval rank, transposed-rank, normalized inverse rank (NIR), and discrimination score (30, 32, 36, 47–49). Key contributions of this work include the extension of the PDS to different underlying dissimilarity measures (including wMSE and wPearsonΔPert, rather than the standard unweighted MSE), its systematic use as a meta-metric for evaluating ground-truth signal quality (rather than model quality), and the gene-wise formulation for characterizing signal sparsity across individual genes.

This extension, which we term the *weighted PDS* (wPDS) or to make the underlying metric explicit PDS (wMSE), substantially improves signal sensitivity by ensuring that the distance computation itself is focused on biologically informative genes. See Liu et al. (32) for an analysis of the scale-sensitivity of the unweighted PDS in high dimensions, where uninformative genes dilute signal with noise; gene weighting directly mitigates this issue by concentrating the distance computation on the subspace carrying perturbation-specific information.

### D. Evaluating metric signal

Before benchmarking predictive models, it is critical to verify that the chosen evaluation metric captures sufficient biological signal in the ground-truth data. We assess metric sensitivity along two complementary dimensions.

#### Perturbation Discrimination Signal

When used as a meta-metric (as distinct from its use as a model evaluation metric), the PDS is computed exclusively from ground-truth samples to quantify the strength of perturbation-specific signal captured by a given dissimilarity measure *d*. A PDS near 0 or lower indicates that the chosen metric completely fails to resolve perturbation identities, implying a limiting signal-to-noise ratio where within-perturbation variance masks between-perturbation differences. We compute the PDS across various bag sizes *n*_bag_, the number of cells present in a query or reference profile, and dissimilarity measures to characterize signal dilution. We also employ the gene-wise form to evaluate signal sparsity.

#### Bound Discrimination Signal

While the PDS measures separation between different perturbations, the Boundary Discrimination Score (BDS) assesses whether a metric can distinguish a perturbation from uninformative background noise. For each perturbation *p*, we partition the ground-truth cells into two halves: one serves as the evaluation target *X*_*p*_ and the other as the technical replicate (positive base-line 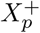). We define a negative baseline 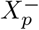 (here, control samples) representing an uninformative prediction. The empirical bounds are 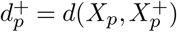(positive bound) and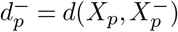 (negative bound). The BDS is the fraction of perturbations where the positive bound is strictly smaller:

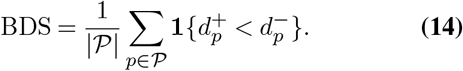

A BDS of 1.0 indicates perfect discrimination of the technical replicate from the uninformative control across all perturbations; a low BDS reveals that the metric lacks the sensitivity to resolve biological signal from background noise (or that the data itself does not contain identifiable signal). Because both the BDS and PDS are derived from a single data partition, we extend them to rigorous permutation tests in Supplementary Note 5 to assess the robustness of these findings across different data splits.

### E. Signal exposure techniques

Evaluating metrics across the entire transcriptome can result in signal dilution, where sparse perturbation-specific effects are masked by the large number of uninformative genes (Supplementary Note 3; see also the empirical characterization in Results). We identify several techniques to expose the biological signal of interest.

#### DEG-based weighting

Following (29, 36), we assign non-negative weights {*w*_*p,g*_} to genes based on their differential expression significance. Specifically, for each perturbation *p, w*_*p,g*_ is derived from the test statistics of gene *g*’s differential expression relative to other perturbed cells in the dataset. This continuous weighting reduces the effective evaluation dimensionality smoothly, rather than through a hard cutoff that would discard genes just below an arbitrary threshold. We extend this approach in two ways. First, we compute DEG weights across the entire dataset (both training and testing partitions), rather than exclusively from the training set. Since these weights serve only for evaluation, this yields a test-set-dependent metric that we consider appropriate. See Supplementary Note 8 for a model rankings with training-set-only weights as computed in (36). Second, we apply continuous gene weighting to the Pearson Delta metric and to the PDS, which to our knowledge has not been done previously.

Concretely, weights are derived as follows. We generate 100 pseudo-bulked dataset mean references by computing the mean expression of |*S*_*k*_| = 100 randomly sampled non-control cell subsets: 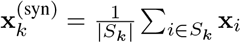. A Welch’s *t*-test, implemented via the python library scanpy (50), is then performed between the perturbation cells and these synthetic controls to yield a raw test statistic *s*_*p,g*_ for each gene *g*. The absolute scores *a*_*p,g*_ = |*s*_*p,g*_| are min-max normalized across genes for each perturbation and squared to emphasize the most strongly differentially expressed genes:

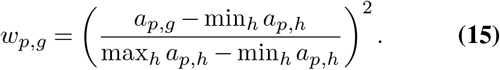

#### DEG filtering

An alternative approach is to restrict evaluation to the top *n* differentially expressed genes per perturbation, a standard strategy that discards uninformative dimensions entirely and introduces no weighting of kept dimensions. In Fig. 5a we compare the PDS signal at varying numbers of selected genes for perturbation-specific DEG filtering and for the same selection of genes for all perturbations, based on mean-aggregated weights. We also compare DEG-filtered and weighted metrics in Supplementary Note 7.

#### Pathway-focused evaluation

Following (37, 38), evaluation can be restricted to genes mapping to specific annotated biological pathways, providing a targeted biological filter. While we do not employ pathway-focused evaluation in the present study, it represents a natural way to incorporate biological prior knowledge for dimensionality reduction and signal exposure, and to obtain interpretable, pathway-specific assessments of model performance.

#### Delta metrics

As proposed in (11, 47), expression values are centered relative to a negative baseline *µ*^−^ before computing dissimilarities, isolating perturbation-specific log-fold changes from the generic cellular state. This centering can also reduce the effective intrinsic dimensionality of the evaluation space: if perturbation-induced changes are confined to a lower-dimensional subspace, subtracting the shared baseline removes variation along uninformative dimensions and concentrates the metric on the subspace where perturbation effects reside.

### F. Effective gene number

Because signal exposure techniques alter the effective evaluation dimensionality, we consistently report the effective gene number *n*_eff_, which represents the approximate number of genes that meaningfully contribute to the evaluation after weighting; intuitively, the “equivalent number” of equally weighted genes. For hard-filtering approaches (DEG or pathway filtering), *n*_eff_ is simply the count of retained genes. For continuous weighting, we compute

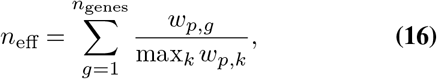

where weights are rescaled such that the maximum equals 1. Reporting *n*_eff_ prevents the evaluation from collapsing onto a trivially small gene subset and enables fair comparison across signal exposure techniques. The weighting scheme and resulting *n*_eff_ can themselves be optimized as shown in Fig. 5.

### G. Metric calibration

To transform raw metric values into interpretable quantities, we introduce perturbation-wise metric calibration relative to the empirical bounds established above. For a model prediction 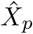 with raw dissimilarity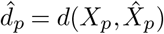, the calibrated score is

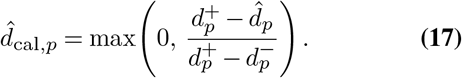

This maps the positive baseline (technical duplicate) to 0 and the negative baseline (uninformative predictor) to 1, with values exceeding 1 indicating worse-than-uninformative performance. The score is clipped at 0 from below to cap predictions that nominally surpass the technical replicate. Perturbations where 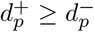 are excluded, as the calibration is not meaningful when the positive bound does not exceed the negative bound. This exclusion criterion is directly related to the BDS: a low BDS indicates that many perturbations fail this condition, signaling that the chosen metric lacks sufficient sensitivity for calibrated evaluation. Practitioners should therefore verify the BDS before interpreting calibrated scores; one may additionally require a minimum separation between the upper and lower bounds as a stricter inclusion criterion, focusing evaluation on perturbations with sufficiently strong discriminability. The choice of bounds is flexible: in the absence of a clean control, the dataset mean or another well-defined reference can be used as the negative baseline, and the calibration interpretation adapts accordingly.

This calibration standardizes performance scales across perturbations and datasets, enabling direct comparison of model improvements relative to the achievable optimum. We note that the Dynamic Range Fraction (DRF) introduced by Miller et al. (36) is trivially saturated under this calibration. The DRF is defined as 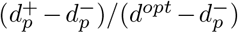 for distance-based metrics; substituting the calibrated metric 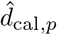, the positive and negative bounds become 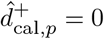and 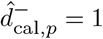 and the optimal metric value *d*^*opt*^ = 0 by construction, yielding DRF = (0 −1)*/*(0 −1) = 1 for every valid perturbation. This raises fundamental questions about the suitability of the DRF as a standalone meta-metric for assessing the intrinsic quality of evaluation metrics. We show in Supplementary Note 4 that the calibrated *R*^2^ Delta Perturbation and calibrated MSE are algebraically equivalent, explaining their identical BDS values.

### H. Datasets

Our benchmarking analysis uses seven perturbational transcriptomics datasets spanning two prediction tasks. For the combinatorial (double) perturbation prediction task, we evaluate on three datasets: norman19 (5), wessels23 (7), and replogle20 (6). For the single perturbation prediction task, we evaluate on four datasets: replogle22k562 (3), nadig25hpeg2 (4), frangieh21 (2), and adamson16 (1). Six of these seven datasets were previously benchmarked by Miller et al. (36); we introduce the replogle20 dataset as a new combinatorial perturbation benchmark, making this the first study to include it for double perturbation prediction evaluation.

The replogle20 dataset originates from Replogle et al. (6), who developed direct-capture Perturb-seq and demonstrated its utility for investigating genetic interactions in K562 cells using both CRISPRi and CRISPRa. The combinatorial screen targets pairs of genes involved in cholesterol biogenesis and DNA repair pathways, providing a biologically focused complement to the broader screens of norman19 and wessels23. The screen contains only 40 double perturbations, of which 10 are available for training under our held-out evaluation split, making replogle20 the smallest combinatorial dataset in our benchmark. We preprocessed replogle20 using the same pipeline applied to norman19.

Table 1 summarizes the key characteristics of all seven datasets, including cell counts, gene numbers, perturbation counts, cell type, and CRISPR modality.

**Table 1.** Overview of perturbational transcriptomics datasets used in this study. Cell, gene, and perturbation counts reflect the fully preprocessed datasets. Single and double perturbation counts are listed separately where applicable.

| Dataset | Cell type | Modality | Cells | Genes | Single perts | Double perts |
| --- | --- | --- | --- | --- | --- | --- |
| *Combinatorial perturbation prediction* |  |  |  |  |  |  |
| norman19 | K562 | CRISPRa | 88,628 | 8,221 | 100 | 124 |
| wessels23 | Monocytes | CRISPRi | 22,794 | 8,210 | 28 | 157 |
| replogle20 | K562 | CRISPRi | 26,938 | 8,212 | 42 | 40 |
| *Single perturbation prediction* |  |  |  |  |  |  |
| replogle22k562 | K562 | CRISPRi | 237,024 | 8,370 | 2,045 | — |
| nadig25hpeg2 | HepG2 | CRISPRi | 109,183 | 8,746 | 2,323 | — |
| frangieh21 | Melanoma | CRISPR-Cas9 | 47,571 | 8,319 | 241 | — |
| adamson16 | K562 | CRISPRi | 55,021 | 8,247 | 91 | — |

### I. Data preprocessing

We adopt the preprocessing pipeline of Miller et al. (36) without modification. For each dataset we filter out perturbations with fewer than 12 measured cells, cells with fewer than 200 expressed genes, and genes expressed in fewer than 3 cells. Expression counts are library-size normalized to a target sum of 10,000 and log1p-transformed (natural logarithm) using scanpy (50). To balance the datasets, a maximum number of cells per perturbation is enforced at the mean cell count per perturbation after filtering. The gene set for each dataset comprises the union of the top 8,192 highly variable genes (identified via `scanpy.pp.highly_variable_genes`) and all perturbed genes. For the replogle20 dataset, we applied the identical preprocessing pipeline described above, consistent with the treatment of norman19.

### J. Evaluation setup

We follow the cross-validation protocol of Miller et al. (36). For the single perturbation prediction task, we perform 5-fold cross-validation with a 60/20/20% train/validation/test split in each fold (perturbations are split, not cells). For the combinatorial perturbation prediction task, we perform 2-fold cross-validation over the double perturbation conditions: all single perturbations are retained in the training set, while double perturbations are randomly divided into two equal halves. In each fold, one half is further split equally into training and validation subsets, so that 25% of all double perturbations are used for training and 25% for validation, with the remaining 50% held out for testing.

Our evaluation codebase is derived from the publicly available benchmark repository of Miller et al. (36) (`github`). We did not modify the data preprocessing, train-validation-test splitting, or model training procedures. Our contributions consist of (i) additional evaluation metrics (wPearsonΔPert, PDS with weighted metrics, and calibrated variants), (ii) a new DEG weighting scheme (see Signal exposure techniques), (iii) the perturbation discrimination analysis infrastructure, (iv) the inclusion of the replogle20 dataset for combinatorial perturbation evaluation and (v) the addition of another state-of-the-art model PDAE.

### K. Models

We benchmark four deep learning models previously evaluated by Miller et al. (36): GEARS (11), scGPT (19), scLambda (34), and PRESAGE (21). In addition, we include PDAE (35), a perturbation prediction model with theoretical identifiability guarantees not evaluated in the original benchmark. All models were trained using their default hyperparameters within the model-specific Docker containers provided by the Miller et al. codebase, with no modifications to training procedures. PDAE was trained with hyperparameters recommended by the authors.

#### Correction for incomplete gene predictions in scGPT

scGPT does not always produce predictions for the full gene set; a subset of genes may be absent from the model’s output. In the original benchmark of Miller et al. (36), this was handled by restricting metric evaluation to the subset of genes for which scGPT produced predictions. However, this approach introduces systematic bias: if genes that are difficult to predict or that carry strong perturbation signal (e.g., highly variable genes or DEGs) are disproportionately excluded, the resulting metric values are computed over an easier, less informative gene subset. This effect is particularly pronounced for weighted metrics such as wMSE, where the excluded genes may receive high DEG-based weights, thereby artificially deflating scGPT’s error relative to models that predict the full transcriptome. We resolved this by imputing all missing gene predictions with the uninformative dataset mean baseline *µ*^all^, ensuring that all models are evaluated on an identical gene set. This correction accounts for the strong wMSE performance of scGPT reported in Miller et al. (36) and places it on equal footing with models that predict all genes.

## Code Availability

The code to reproduce the results presented here is available at https://github.com/michavol/sbb-perturbation-benchmark.

## Acknowledgements

The authors thank H. Miller for technical assistance with code repositories, L. Rabuzin for feedback on methodology, and M. Ulmer and J. Ketterer for discussions regarding evaluation practices and perturbation prediction models. M. Vollenweider gratefully acknowledges financial support from the Swiss National Science Foundation (grant no. 214865).

## Author Contributions

M.Vollenweider conceived the study, designed and performed the experiments, developed the methodology and software, and conducted all data analyses. P.Bühlmann provided project guidance and supervised the research. M. Vollenweider wrote the original draft of the manuscript, and both authors contributed to the final interpretation of the results and to editing the manuscript.

## Declaration of Interest

The authors declare no competing interests.

|  |  |  |  |
| --- | --- | --- | --- |
| PROJECT DETAILS | PROJECT NAME<br>SBB Principles | DOMAIN | KEY APPLICATION |
| CONTACT(S) | NAME(S)<br>Michael Vollenweider | EMAIL(S) | AFFILIATION(S)<br>ETH Zurich, Seminar of Statistics |
| MODEL(S) | MODEL NAME(S)<br>Gemini 3, Claude 4.6 Opus, Claude 4.7 Opus, Composer 2 |  |  |
| LITERATURE REVIEW | FINDING LITERATURE<br>Gemini 3 | FINDING EXAMPLES FROM KNOWN LITERATURE OR ADDING LITERATURE FOR EXISTING STATEMENTS | COMPARING LITERATURE<br>Gemini 3 |
| WRITING | GENERATING NEW TEXT BASED ON INSTRUCTIONS<br>Claude 4.6 Opus, Claude 4.7 Opus | ASSISTING IN IMPROVING OWN CONTENT OR PARAPHRASING RELATED WORK<br>Claude 4.6 Opus | PUTTING OTHER WORKS IN PERSPECTIVE |
| CODING | GENERATING NEW CODE BASED ON DESCRIPTIONS OR EXISTING CODE<br>Claude 4.6 Opus, Composer 2 | REFACTORING AND OPTIMIZING EXISTING CODE<br>Claude 4.6 Opus, Composer 2 | COMPARING ASPECTS OF EXISTING CODE |
| ETHICS | WHY DID WE USE AI FOR THIS PROJECT?<br>Efficiency | WHAT STEPS ARE WE TAKING TO MITIGATE ERRORS OF AI?<br>Double-checking sources and generated code | WHAT STEPS ARE WE TAKING TO MINIMIZE THE CHANCE OF HARM OR INAPPROPRIATE USE OF AI? |
| THE CORRESPONDING AUTHORS VERIFY AND AGREE WITH THE MODIFICATIONS OR GENERATIONS OF THEIR USED AI-GENERATED CONTENT |  |  |  |

AI Usage Card v2.0
https://ai-cards.org

## Supplementary Note 1: Calibrated performance heatmaps across datasets

**Fig. 6.**
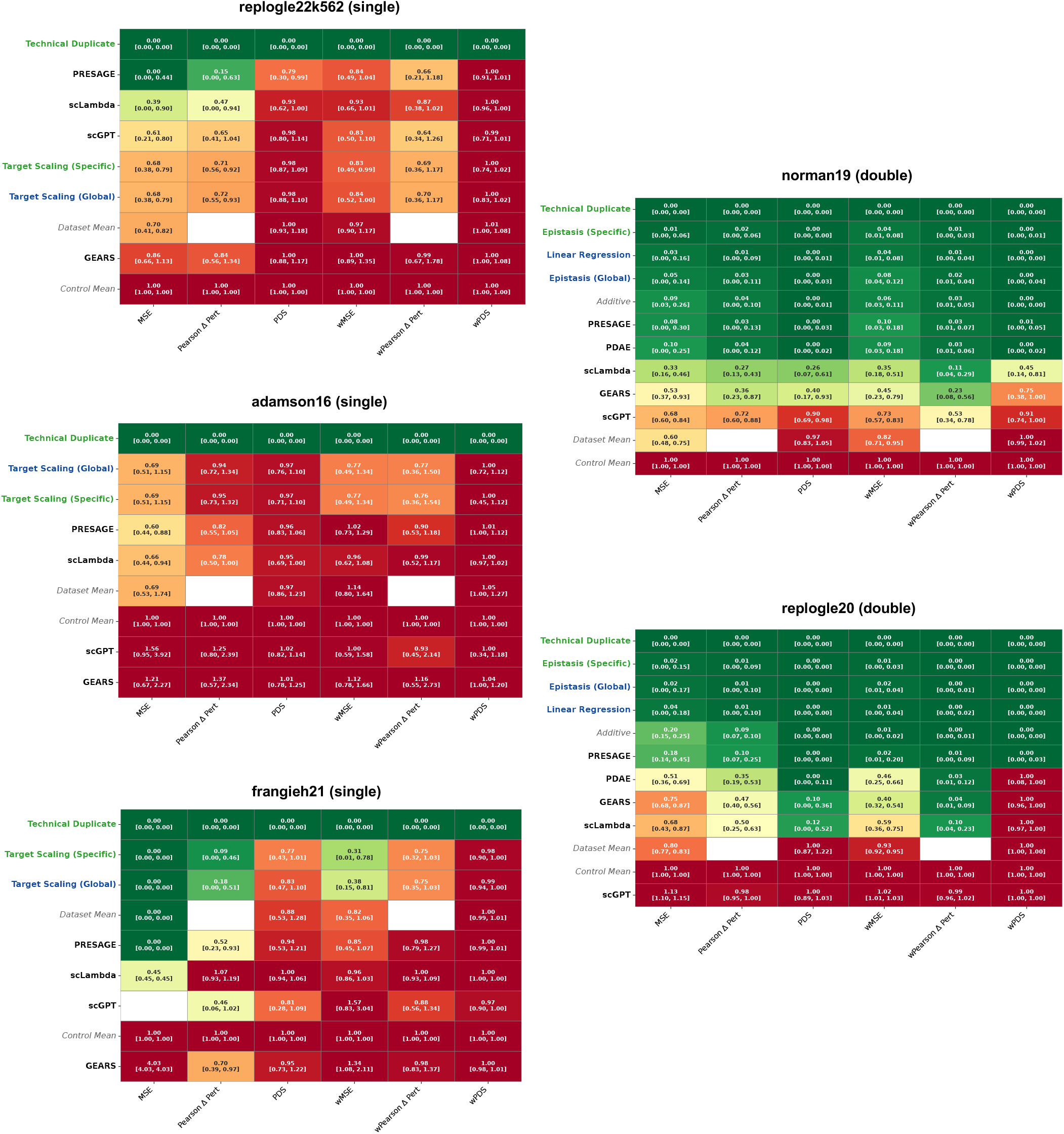
Calibrated performance heatmaps. Calibrated performance heatmaps for single and double perturbation prediction; values represent median calibrated scores (0 = technical duplicate, 1 = uninformative control) and brackets indicate the 25th and 75th percentile. For frangieh21 the entry for scGPT and MSE is blank because in at least one test fold the BDS is 0, so calibration is not possible for any perturbation in this fold. Also see Fig. 4a.

## Supplementary Note 2: Detailed results for wessels23 & nadig25hpeg2

**Fig. 7.**
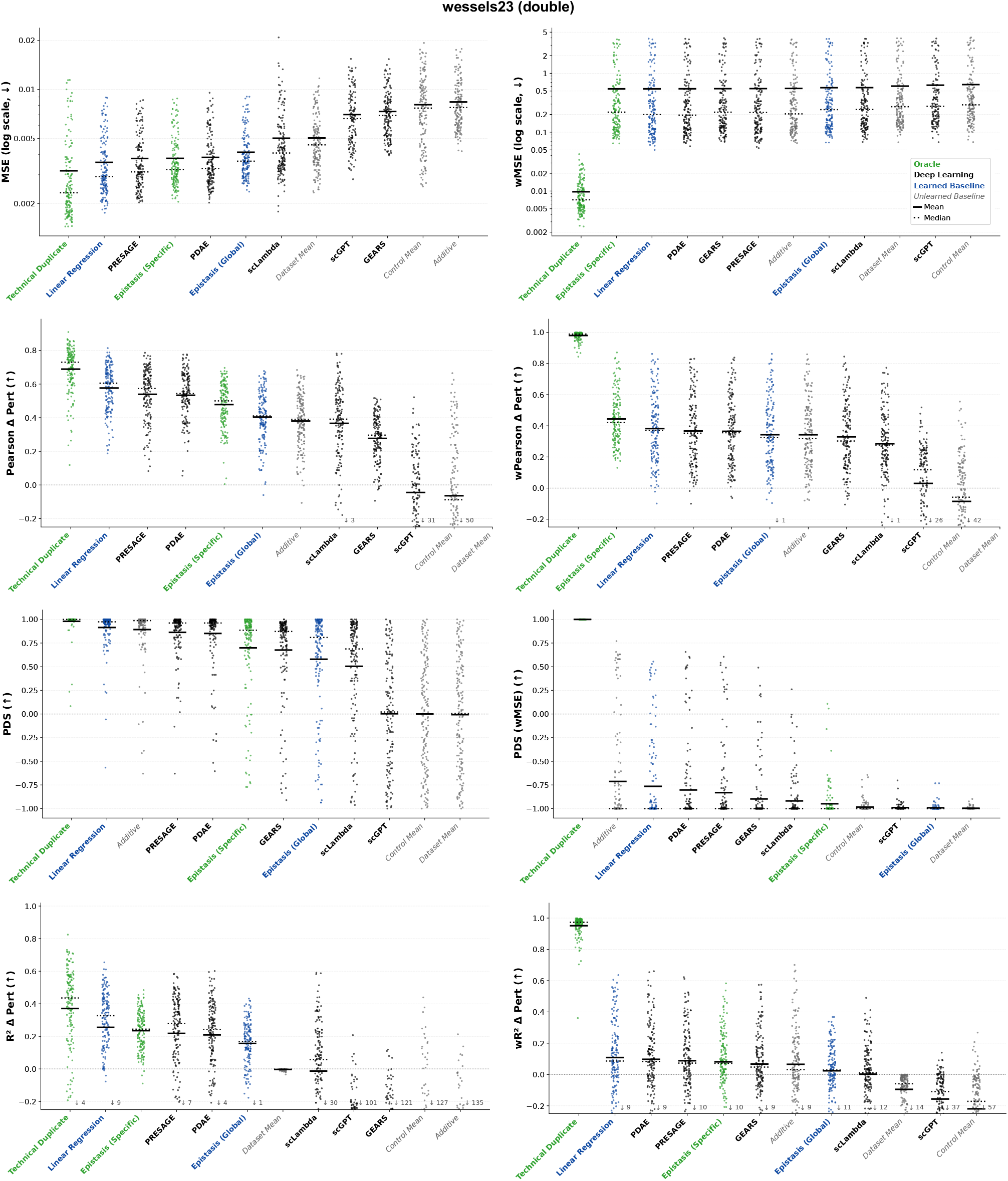
Detailed results for wessels23. Per-perturbation metric values across models. Sorted by mean across perturbations.

**Fig. 8.**
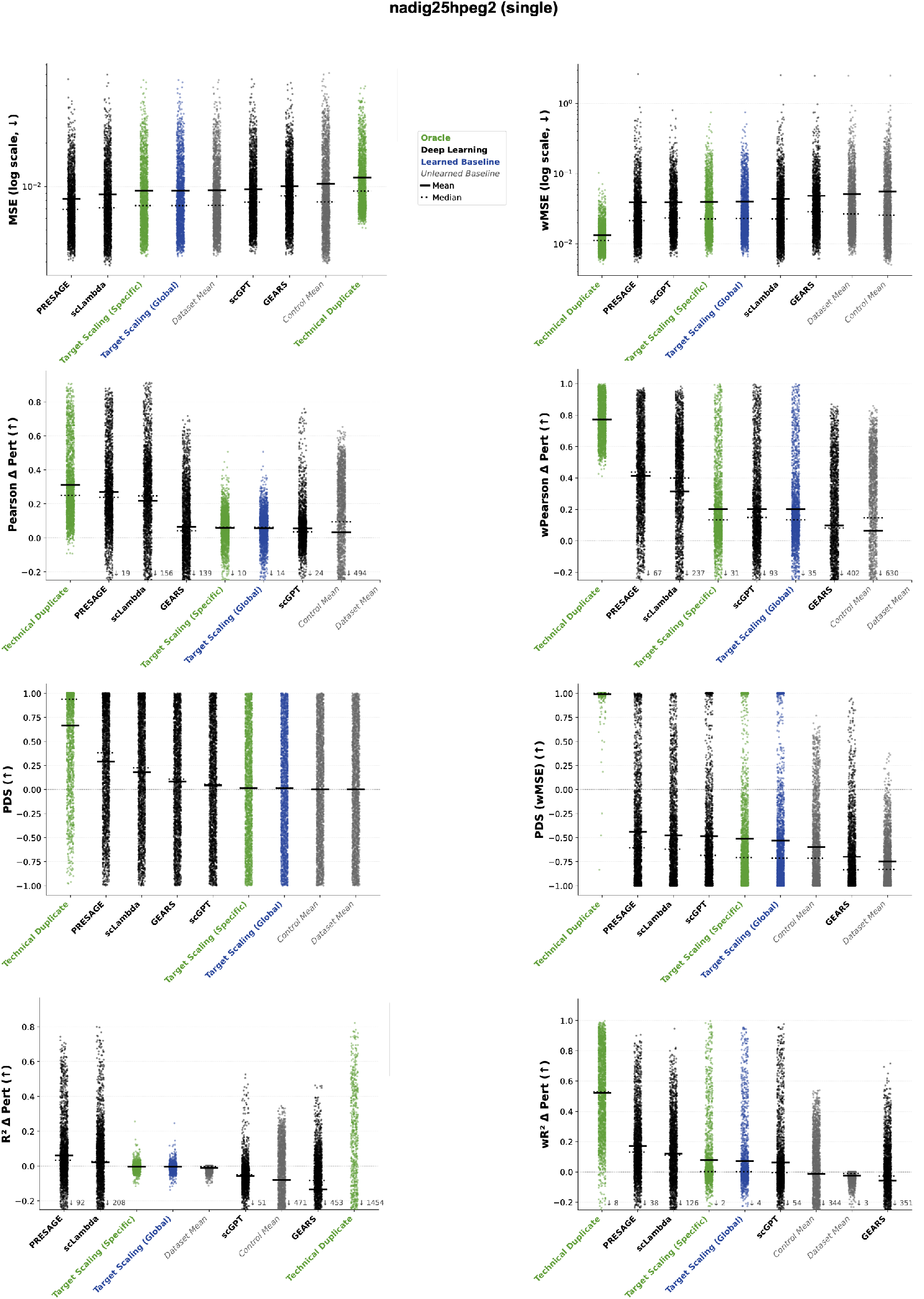
Detailed results for nadig25hpeg2. Per-perturbation metric values across models. Sorted by mean across perturbations.

## Supplementary Note 3: Mathematical intuition for signal dilution

We formalize how high-dimensional noise inevitably dilutes sparse biological signals during evaluation and how the signal exposure techniques discussed in the main text systematically increase the signal-to-noise ratio.

### A. Distance concentration in high dimensions

Consider two independent random transcriptomic vectors **x, y** ∈ ℝ^*n*^ with independent, standardized compon ents *x*_*g*_, *y*_*g*_ ~ 𝒩 (0, 1). By the law of large numbers, the squared norms concentrate around the ambient dimension: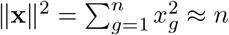. The inner product **x** · **y** = Σ _*g*_ *x*_*g*_*y*_*g*_ is a sum of *n* independent zero-mean, unit-variance terms, so

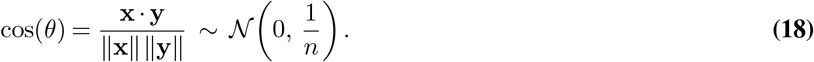

As *n* → ∞, the cosine similarity concentrates at zero: random high-dimensional vectors are asymptotically orthogonal. The squared Euclidean distance consequently concentrates around a deterministic constant:

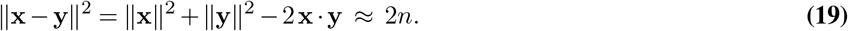

This concentration phenomenon is the root cause of signal dilution: when a distance metric is dominated by noise dimensions, all pairwise distances collapse to the same value, making biologically distinct conditions indistinguishable.

### B. Signal dilution in mixed signal–noise subspaces

Biological expression vectors are not purely random. Suppose each vector contains a signal subspace of *n*_*S*_ genes exhibiting pairwise correlation *ρ >* 0 and a noise subspace of *n*_*N*_ = *n* − *n*_*S*_ genes with independent entries. Define the signal fraction *α* = *n*_*S*_*/n*. The expected inner product decomposes as

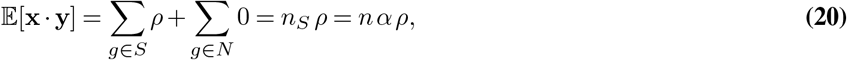

yielding, for the observable cosine similarity and squared distance,

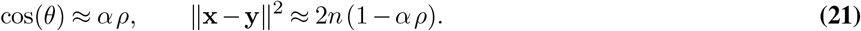

When *α* is small, as is typically the case for perturbation-specific effects, the cosine similarity shrinks toward zero and the distance approaches the pure-noise limit 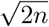, masking genuine biological similarity.

### C. Perturbation-specific signal-to-noise ratio

In the context of perturbation evaluation, the deterministic biological state of a cell under perturbation *p* can be decomposed as **s**_*p*_ = **s**_generic_ + **s**_pert,*p*_, where **s**_generic_ captures the shared cellular state and **s**_pert,*p*_ the perturbation-specific shift. A single observed cell is 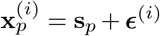, where ***ϵ***^(*i*)^ is independent high-dimensional noise with ∥***ϵ***^(*i*)^∥^2^ ≈ *n*.

For the PDS to yield a high score, the within-perturbation distance (between cells sharing the same **s**_pert,*p*_) must be distinguishably smaller than the cross-perturbation distance (between cells with different perturbation-specific shifts). This requires sensitivity specifically to **s**_pert,*p*_, motivating the perturbation-specific signal-to-noise ratio:

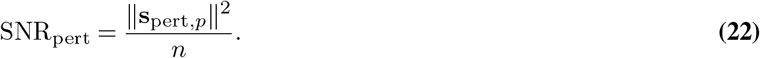

Empirically, **s**_pert,*p*_ is sparse: it is confined to a small number of genes *n*_pert_ ≪ *n*, so the signal fraction *α*_pert_ = *n*_pert_*/n* is tiny. As a result, both inter- and intra-group distances collapse toward the noise limit 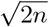, rendering perturbations indistinguishable under standard metrics.

### D. Recovering signal through pseudobulking and weighting

Two complementary strategies increase SNR_pert_.

#### Pseudobulking (reducing the noise floor)

Averaging *k* cells from the same perturbation yields 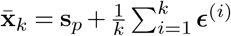, with residual noise magnitude 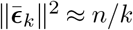. The effective SNR scales linearly with cell count:

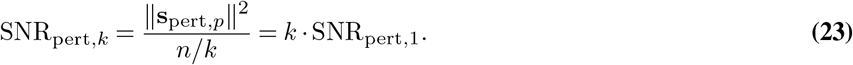

As *k* increases, the noise floor drops and distance metrics begin to resolve **s**_pert,*p*_.

#### Gene weighting and filtering (increasing the signal fraction)

Alternatively, one can increase *α*_pert_ directly by reducing the dimensionality. DEG-based gene weighting, pathway filtering, and delta centering all suppress the *n*_*N*_ noise dimensions, concentrating the metric on the *n*_*S*_ signal dimensions and preventing distance concentration. This explains the empirical observation (Fig. 4c) that weighted metrics maintain high PDS across the full gene set even at low cell counts, while unweighted MSE suffers severe dilution.

## Supplementary Note 4: Equivalence of calibrated 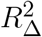 and calibrated MSE

We show that the calibrated *R*^2^ Delta Perturbation metric and the calibrated MSE reduce to the same expression, explaining why they yield identical BDS values and calibrated scores in the heatmaps. Recall from Methods that for a perturbation *p* with observed mean *µ*_*p*_, the MSE of any predictor 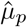is 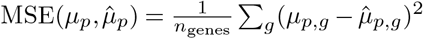. Calibration maps the raw MSE into the interval defined by the positive bound (technical duplicate, MSE_pos_) and negative bound (uninformative control, MSE_neg_):

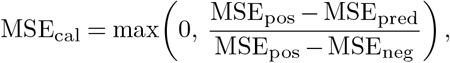

where the max(0, ·) clips predictors that perform worse than the negative bound to zero. For the argument below we omit this clipping, since it is applied identically to both metrics and therefore does not affect their equivalence. The 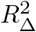 metric is defined on expression changes relative to the negative baseline *µ*^−^. Writing 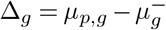 and 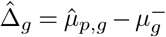, the metric is

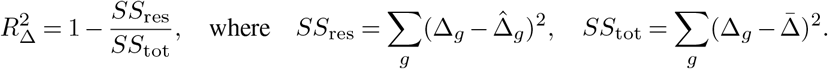

The key observation is that the negative baseline cancels in the residual:

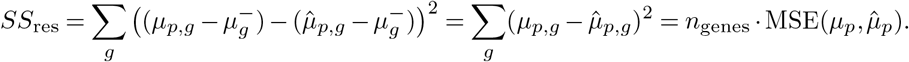

Thus *SS*_res_ is proportional to MSE regardless of the choice of *µ*^−^. The calibrated 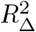 is

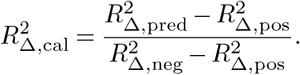

Substituting 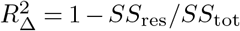 into each term, we note that *SS*_tot_ depends only on the ground-truth expression changes Δ_*g*_ and their mean 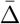, so it takes the same value whether we evaluate the model prediction, the technical duplicate, or the uninformative control. We get:

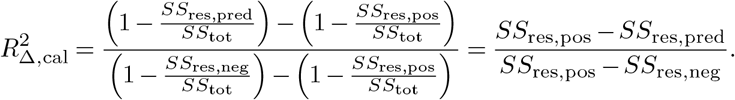

Since each *SS*_res_ term equals *n*_genes_ · MSE, the constant *n*_genes_ cancels, giving

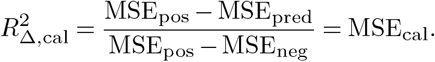

The calibrated 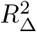 is therefore algebraically identical to the calibrated MSE for every perturbation.

## Supplementary Note 5: Metric evaluation via permutation testing

The BDS and PDS provide intuitive point estimates of biological signal strength, but they are derived from a single partition of the cells into query and reference bags and therefore lack formal null distributions. To robustly quantify the statistical significance of these scores and account for sampling variance, we extend both into permutation-based testing procedures. These are instances of multivariate two-sample permutation tests (52), a well-established family of nonparametric methods that construct null distributions by randomly reassigning group labels while preserving the overall data structure. Because permutation tests require no distributional assumptions on the high-dimensional expression vectors, they are naturally suited to single-cell transcriptomic data. The resulting *p*-values provide exact finite-sample type I error control under the null hypothesis of exchangeability. Beyond their role in validating evaluation metrics, these tests are independently useful for characterizing the data itself. The PDS-based test identifies which perturbations produce statistically detectable transcriptomic signatures under a given dissimilarity measure, while the BDS-based test identifies which perturbations are distinguishable from the control background. This information can guide experimental quality control, help select appropriate evaluation metrics, and inform the design of perturbation screens by flagging conditions with insufficient signal prior to any model benchmarking.

### PDS-based testing

To test whether a perturbation’s transcriptomic signature is significantly distinct from the rest of the library, we evaluate the rank of its within-perturbation dissimilarity among all cross-perturbation dissimilarities. For a given perturbation *p*∈ 𝒫, let 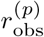 be the ordinal rank of *d*(*Q*_*p*_, *R*_*p*_) when pooled with all cross-perturbation dissimilarities *d*(*Q*_*p*_, *R*_*q*_) for *q* ≠ *p*, where *Q*_*p*_ and *R*_*p*_ denote the query and reference bags as defined in Methods (Eq.13). We construct a null distribution by randomly permuting the perturbation labels across all 2|𝒫 | half-bags *M* times, preserving the bag structure while breaking the association between bags and perturbation identities. For each permutation *m*, we recompute the rank 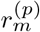 under the permuted labels. The empirical *p*-value is the fraction of permutations where the rank is as small or smaller than the observed rank:

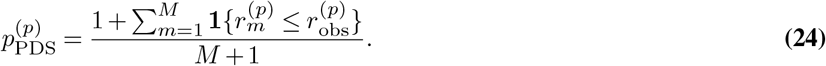

A small 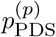 indicates that perturbation *p* produces a transcriptomic signature that is significantly more self-similar than expected under random label assignment, confirming that the dissimilarity measure *d* can resolve this perturbation’s identity. Ideally, one would also permute over the splitting into *Q* and *R* sets, but this incurs a significantly higher computational burden.

### BDS-based testing

The BDS tests whether the distance between technical replicates (positive bound) is smaller than the distance to control cells (negative bound). For each perturbation *p*, we compute the observed difference 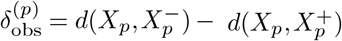, where *X*_*p*_ is the evaluation target, 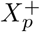 is the technical replicate, and 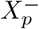 is the control baseline, following the notation in Methods. Under the null hypothesis that perturbed and control cells are exchangeable, this difference should not be systematically positive. We construct a null distribution by pooling the cells of perturbation *p* with the control cells and randomly reassigning their condition labels *M* times, recomputing 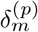 for each permutation. The empirical *p*-value is:

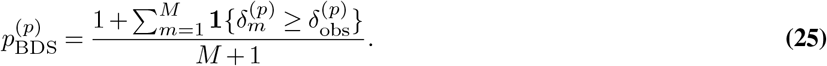

This is structurally analogous to a two-sample permutation test based on a distance statistic (53), where the test statistic is the difference between inter-group and intra-group dissimilarities. A small 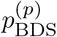 confirms that the perturbation’s effect is reliably distinguishable from the control background under the chosen metric.

### Aggregation across perturbations

Both procedures yield a *p*-value for every individual perturbation *p*. To summarize the overall sensitivity of a metric *d* across the perturbation library, we apply the Benjamini-Hochberg procedure to control the false discovery rate (FDR) and report the fraction of perturbations with statistically significant discrimination. This transforms the single-split BDS and PDS into robust, distribution-backed assessments of metric adequacy, complementing the point estimates reported in the main text.

### Validity under weighted metrics

When the dissimilarity *d* is a weighted metric such as wMSE or wPearsonΔPert, the DEG-based weights are computed from the full dataset, including the test perturbations whose significance is being assessed. This introduces a dependency between the test statistic and the data used to construct it, which could in principle inflate significance by biasing the dissimilarity toward dimensions where signal happens to be present. However, the permutation procedure partially mitigates this concern: because the same fixed weights are applied under both the observed and permuted label assignments, the null distribution is constructed under the same weighting scheme, so the comparison remains internally consistent. The weights do not change across permutations; only the label assignments do. A stricter approach would recompute weights within each permutation, but this is computationally prohibitive and would also alter the metric being tested. We therefore interpret the resulting *p*-values as valid assessments of whether the observed signal exceeds what would arise by chance under the given (fixed) weighting scheme, while acknowledging that the choice of weighting scheme itself is data-dependent. For unweighted metrics, this concern does not arise.

## Supplementary Note 6: Interpretation of PDS as AUROC

The per-perturbation PDS score *s*_*p*_ defined in Methods can be formally interpreted as a rescaled area under the receiver operating characteristic curve (AUROC) for a distance-based binary classification task. This connection follows from the classical equivalence between the AUROC and the Mann-Whitney *U* statistic (54, 55) and provides both an intuitive probabilistic interpretation and access to the well-developed statistical theory surrounding the AUROC.

### Binary classification setup

Consider a fixed perturbation *p* with query profile *Q*_*p*_. For each other perturbation *q* ∈\ 𝒫{*p*}, we observe a cross-perturbation dissimilarity *d*(*Q*_*p*_, *R*_*q*_). We also observe the within-perturbation dissimilarity *d*(*Q*_*p*_, *R*_*p*_). Define a binary classification problem with a single “positive” instance (the within-perturbation distance *d*(*Q*_*p*_, *R*_*p*_)) and *n*_perts_ − 1 “negative” instances (the cross-perturbation distances {*d*(*Q*_*p*_,*R*_*q*_)}_*q*≠*p*_). The PDS asks whether the positive instance has a smaller value than the negative instances, that is, whether the query is closer to its own reference than to references of other perturbations.

### Connection to AUROC

The AUROC for a binary classifier admits a well-known probabilistic interpretation (54): it equals the probability that a randomly chosen positive instance receives a higher score than a randomly chosen negative instance. In our setting, a “higher score” for the positive class corresponds to a *smaller* dissimilarity (the positive instance should have a lower distance), so we reverse the comparison. Let *D*^+^ = *d*(*Q*_*p*_, *R*_*p*_) denote the positive distance and *D*^−^ denote a cross-perturbation distance *d*(*Q*_*p*_, *R*_*q*_) for a uniformly chosen *q*≠ *p*. The AUROC is the probability

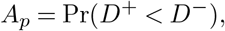

which we estimate empirically by exhaustive comparison of the single positive instance against all *n*_perts_ −1 negative instances:

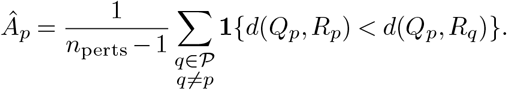

### Rescaling to PDS

Comparing with the PDS definition from Methods, the rank of the within-perturbation distance among all cross-perturbation distances is

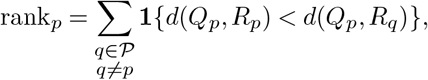

and the per-perturbation PDS score is

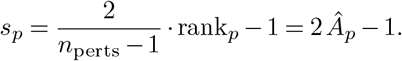

The PDS is therefore a linear rescaling of the AUROC from [0, 1] to [−1, 1], where *s*_*p*_ = 1 corresponds to *Â*_*p*_ = 1 (perfect discrimination), *s*_*p*_ = 0 corresponds to *Â*_*p*_ = 0.5 (chance level), and *s*_*p*_ = −1 corresponds to *Â*_*p*_ = 0 (systematically reversed ordering). The aggregated PDS across all perturbations is

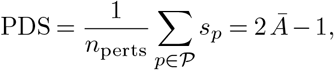

where *Ā* is the mean estimated AUROC across perturbations.

### Interpretation

This equivalence gives the PDS a direct probabilistic reading: *Â*_*p*_ = (*s*_*p*_ + 1)*/*2 estimates the probability that a randomly chosen cross-perturbation distance exceeds the within-perturbation distance for perturbation *p*. A dataset-level PDS of, say, 0.6 corresponds to a mean AUROC of 0.8, meaning that on average, there is an estimated 80% probability that the dissimilarity measure correctly identifies a perturbation’s own replicate as more similar than a randomly chosen other perturbation. This probabilistic framing makes the PDS directly comparable to classification performance metrics used throughout machine learning, and connects it to the extensive statistical theory available for the Mann-Whitney *U* statistic, including variance estimates, confidence intervals, and asymptotic normality results (54).

## Supplementary Note 7: Extended metric sensitivity comparison

This supplementary note extends the metric sensitivity analysis from the main text by comparing metric variants along four axes: (i) DEG-filtered (top-*n*) versus continuously weighted metrics, (ii) distributional single-cell distances versus mean-based metrics, (iii) DEG weights computed against true control cells versus synthetic controls, and (iv) the choice of reference mean used for centering in PearsonΔ variants (no centering, control mean *µ*_0_, or perturbation-average mean *µ*^all^). Comparisons are formalized through the permutation testing framework of Supplementary Note 5.

### Permutation testing across metric variants

Figure 9 presents PDS and BDS permutation tests across all seven datasets for a comprehensive set of metric variants, including weighted and unweighted MSE, PearsonΔPert, *R*^2^ΔPert, and top-20 DEG-filtered variants. Two weighting schemes are compared: weights derived from DEG analysis against synthetic controls (syn), as described in Methods, and weights derived against true control cells (vc).

**Fig. 9.**
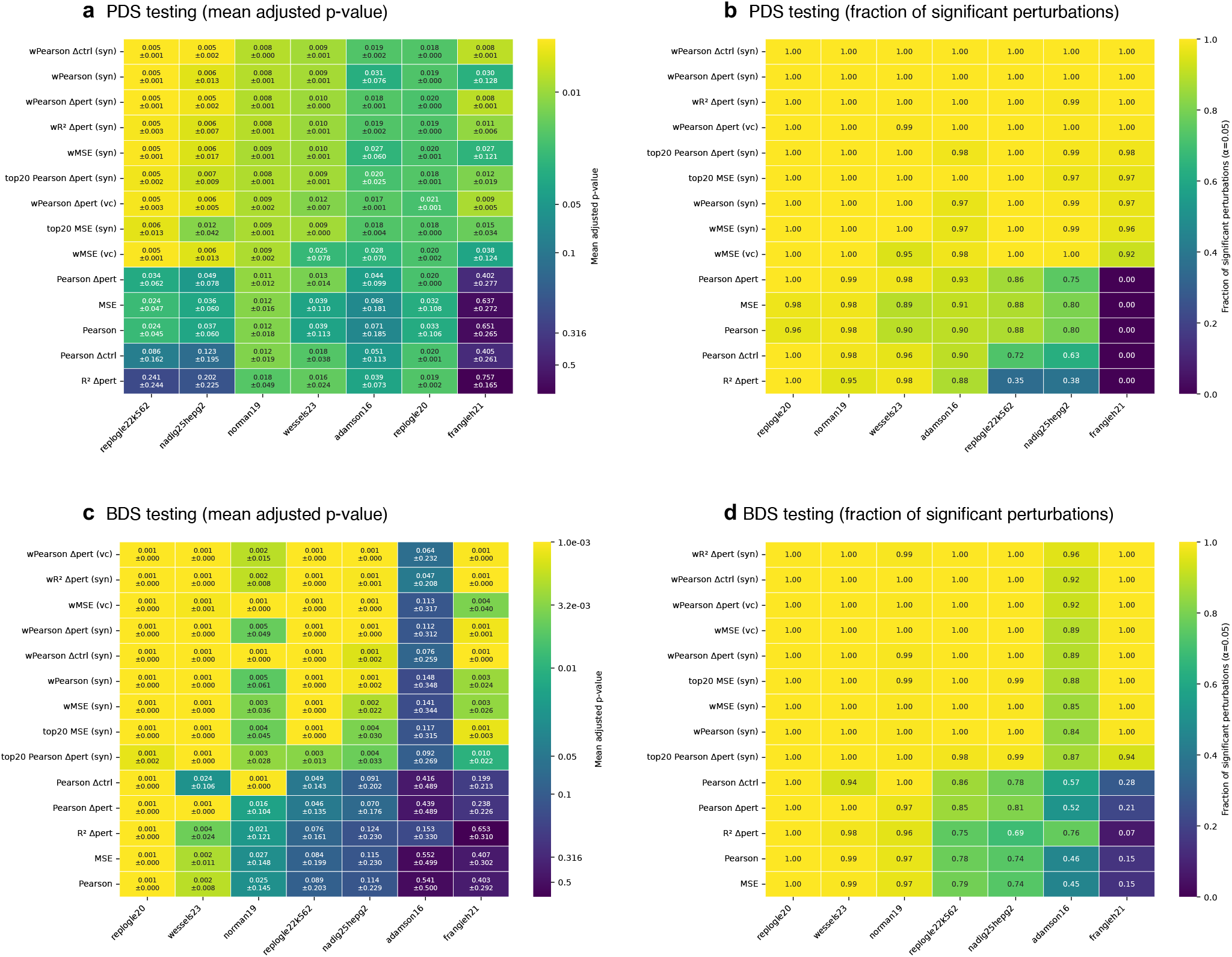
PDS and BDS testing for metric comparison. Permutation tests are performed with 999 permutations as described in Supplementary Note 5. The number of total perturbations considered is limited to 500 perturbations per dataset. With (syn) in brackets, DEG weights are computed as described in Methods. For (vc), DEG weights are derived the same way, but with true control cells as reference, instead of synthetic controls. **a**, Mean adjusted p-value with standard deviation across all perturbations per dataset and per metric. **b**, Fraction of perturbations for which the adjusted p-value for PDS testing is below the significance level of 0.05. **c&d**, Same as for PDS testing but with p-values derived from BDS permutation tests.

The PDS results (Fig. 9a,b) confirm that all weighted and top-*n* filtered variants achieve near-universal significance across datasets (significance fractions ≥0.92). The top-20 variants perform comparably to continuously weighted counterparts, indicating that the precise mechanism of signal concentration matters less than the act of concentrating itself. Among unweighted metrics, all variants fail completely on frangieh21 (significance fraction 0.00), and sensitivity degrades progressively on the large single perturbation datasets. On nadig25hpeg2, MSE and Pearson maintain moderate sensitivity (0.80 each), while PearsonΔCtrl and *R*^2^ΔPert degrade more severely (0.63 and 0.38, respectively). A similar pattern holds on replogle22k562, where MSE and Pearson achieve 0.88 while *R*^2^ΔPert drops to 0.35. On the double perturbation datasets (replogle20, norman19, wessels23), unweighted metrics retain comparatively high sensitivity, though not uniformly so: on wessels23 in particular, MSE and Pearson drop to around 0.89–0.90, and PearsonΔCtrl to 0.96, indicating that even on these datasets weighting offers a non-trivial improvement over unweighted baselines.

The BDS analysis results in (Fig. 9c,d) expose a distinct failure pattern on adamson16 and frangieh21. On adamson16, unweighted MSE and Pearson achieve BDS significance fractions of only 0.45 and 0.46, respectively, while on frangieh21 both drop to 0.15. Notably, the weighted variants restore high BDS significance across all datasets, though adamson16 remains somewhat challenging even under weighting (significance fractions of 0.85–0.96 depending on the weighting scheme). The BDS column ordering differs from the PDS, reflecting dataset-specific difficulty: the datasets most challenging for BDS (adamson16, frangieh21) partially differ from those most challenging for PDS (nadig25hpeg2, frangieh21).

### Pearson correlation reference: control versus perturbation mean

The PearsonΔ metric computes correlation of expression changes relative to a reference mean; we compare three variants: plain Pearson correlation (no centering), PearsonΔCtrl (centering by *µ*_0_, as in Roohani et al. (11)), and PearsonΔPert (centering by *µ*^all^, following Miller et al. (36)). The PDS results (Fig. 9b) reveal that on the large single perturbation datasets nadig25hpeg2 and replogle22k562, PearsonΔPert achieves higher significance fractions than PearsonΔCtrl (0.75 versus 0.63 on nadig25hpeg2; 0.86 versus 0.72 on replogle22k562), consistent with the robustness argument of Viñas-Torné et al. (28). Unexpectedly, however, plain Pearson correlation without any centering matches or outperforms both delta variants on these two datasets (0.80 on nadig25hpeg2 and 0.88 on replogle22k562). This runs counter to the common practice of evaluating perturbation predictions in the Δ space and warrants further investigation into the conditions under which centering is beneficial. On wessels23, conversely, plain Pearson (0.90) is lower than PearsonΔPert (0.98), and for the BDS on wessels23 (Fig. 9d), PearsonΔCtrl yields a lower significance fraction (0.94) than PearsonΔPert (1.00). In all cases, DEG-based weighting renders the choice of reference largely immaterial, as weighted variants of all three metrics achieve near-perfect sensitivity.

### Synthetic versus true control DEG weights

The two weighting schemes yield broadly consistent PDS and BDS results across all datasets (Fig. 9), indicating that the choice of reference population for DEG computation does not materially affect metric sensitivity. The synthetic control approach has the practical advantage of not requiring dedicated control cells, which may be scarce in certain experimental designs, while true control weights may be more straightforward to interpret, as they reflect differential expression against an experimentally defined baseline rather than a computationally constructed one.

### Distributional versus mean-based distance measures

A natural question is whether distributional distance measures that compare full single-cell populations, rather than pseudobulk means, provide superior sensitivity for perturbation discrimination. We consider two such measures. The *energy distance* (53) between two samples 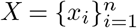 and 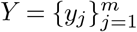 is defined as

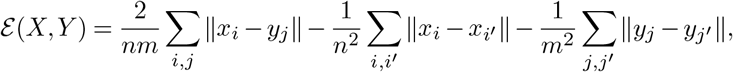

which is zero if and only if *X* and *Y* are drawn from the same distribution (in the population limit). The *Wasserstein distance* (here, the 1D Wasserstein distance used in the gene-wise analysis) between two one-dimensional distributions with CDFs *F*_*X*_ and *F*_*Y*_ is

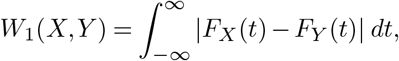

and captures the minimum work required to transport mass from one distribution to the other. Both measures compare full expression distributions rather than their means, and in principle can detect distributional differences (e.g., bimodality, changes in variance) that pseudobulk averaging discards.

Figure 10 addresses this through several complementary analyses. The PDS permutation tests (Fig. 10a,b) compare top-100 energy distance against top-100 MSE and unweighted MSE. Both filtered metrics achieve near-perfect significance fractions across all datasets, with top-100 energy distance performing comparably to top-100 MSE. The gene-wise PDS analysis (Fig. 10d) compares MAE on pseudobulk means, energy distance, and Wasserstein distance: both distributional metrics yield slightly higher per-gene PDS than the mean-based MAE across most datasets, indicating that distributional distances capture additional within-perturbation heterogeneity that pseudobulk averaging discards for the one-dimensional case.

**Fig. 10.**
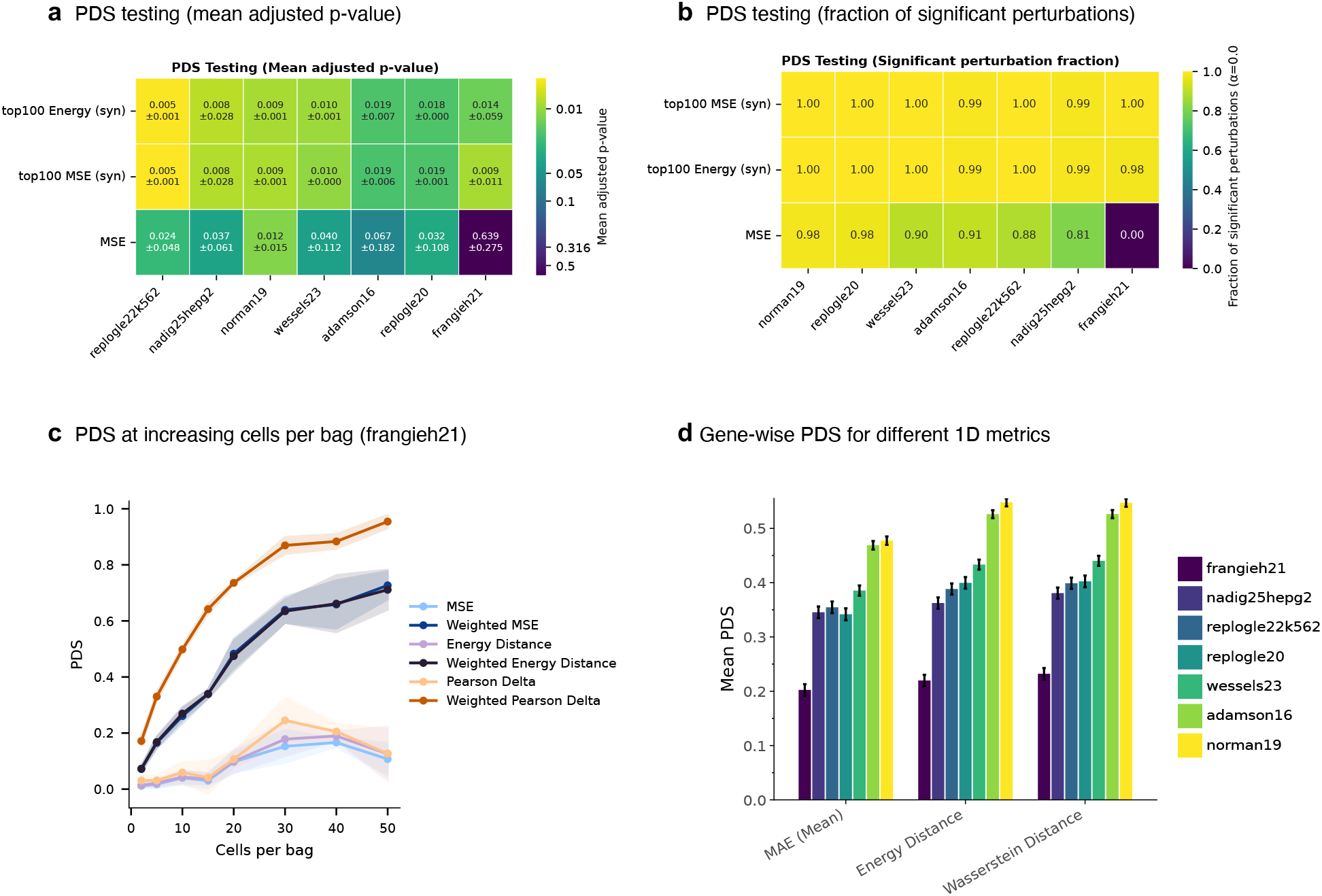
Distributional vs. mean-based metrics. **a**, Fraction of perturbations for which the adjusted p-value for PDS testing is below the significance level of 0.05, comparing top-100 MSE (syn), top-100 energy distance (syn), and unweighted MSE. **b**, Mean adjusted p-value with standard deviation across all perturbations per dataset and per metric. **c**, PDS on technical duplicates with increasing size of reference and query sets on frangieh21, for 50 perturbations and three trials. **d**, Mean gene-wise PDS on the whole dataset for different one-dimensional dissimilarities. MAE (Mean) denotes MAE on the mean vectors of given cell bags.

However, this per-gene advantage does not translate into a substantial improvement at the global level: the PDS permutation test results for top-100 energy distance and top-100 MSE are nearly indistinguishable, and the difference between distributional and mean-based metrics is small relative to the effect of DEG-based weighting or top-*n* filtering. The signal dilution analysis on frangieh21 (Fig. 10c) further illustrates this point: while energy distance achieves marginally higher PDS than MSE at equivalent cell counts, the dominant effect is whether a signal exposure technique (weighting or pseudobulking) is applied at all. Weighted MSE and weighted energy distance both converge to high PDS at moderate cell counts, whereas both unweighted variants suffer from signal dilution. Taken together, these results indicate that the choice between distributional and mean-based distances is secondary to proper signal exposure through gene weighting, and that mean-based metrics, which are computationally much cheaper, provide a reasonable default when combined with DEG-based weighting.

## Supplementary Note 8: Ranking for DEG on training and vs all other cells

**Fig. 11.**
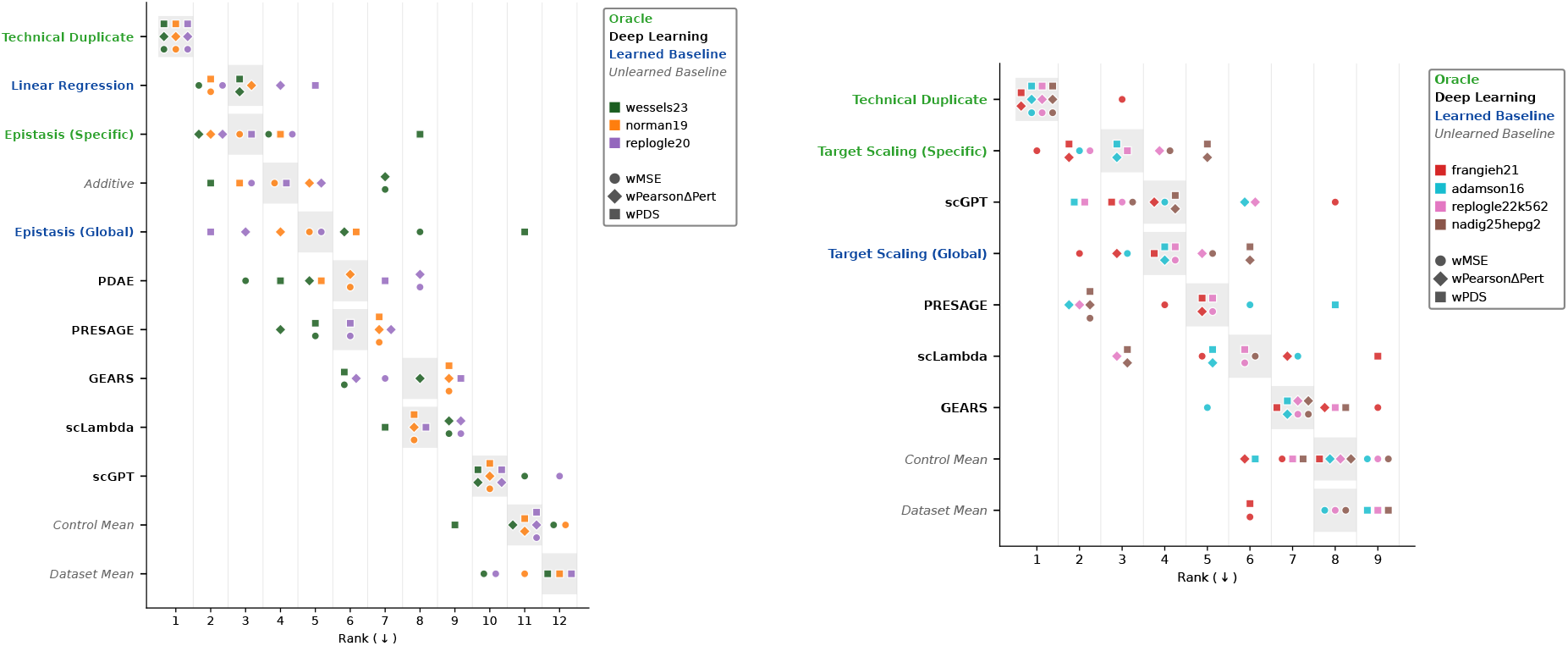
Model ranking for weights derived from DEG analysis on train only. Rankings for single and double perturbation datasets under weighted metrics with weights derived as in (36). Colors distinguish datasets; shapes distinguish metrics. Models are ordered by median rank across all metric-dataset combinations within each panel. The areas shaded in grey mark the overall median rank across datasets and metrics.

